# A bipartite function of ESRRB can integrate signaling over time to balance self-renewal and differentiation

**DOI:** 10.1101/2022.09.20.508291

**Authors:** Teresa E. Knudsen, William Hamilton, Martin Proks, Maria Lykkegaard, Alexander V. Nielsen, Ala Trusina, Joshua M. Brickman

## Abstract

Cooperative DNA binding of transcription factors (TFs) integrates external stimuli and context across tissues and time. Naïve mouse embryonic stem cells are derived from early development and can sustain the pluripotent identity indefinitely. Here we ask whether TFs associated with pluripotency evolved to directly support this state, or if the state emerges from their combinatorial action. NANOG and ESRRB are key pluripotency factors that co-bind DNA. We find that when both factors are expressed, ESRRB supports pluripotency. However, when NANOG is not present, ESRRB supports a bistable culture of cells with an embryo-like primitive endoderm identity ancillary to pluripotency. The stoichiometry between NANOG and ESRRB quantitatively influences differentiation, and *in silico* modeling of bipartite TF activity suggests ESRRB safeguards plasticity in differentiation. Thus, the concerted activity of cooperative TFs can transform their effect to sustain intermediate cell identities and allow *ex vivo* expansion of highly stable stem cell models.

## Introduction

Gene regulation is governed by sequence specific DNA binding proteins, transcription factors (TFs), that integrate diverse stimuli via cooperative binding, where the binding affinity and activity of a TF depends on its partners. In this way, TFs can respond to various environmental cues in a context-specific manner. The exploitation of cooperativity in gene regulation also implies that alterations in one factor’s binding can produce a non-linear switch between transcriptional states. As a result, the combinatorial effect of two TFs can be a distinct output from that generated by one TF on its own. The specificity enabled by cooperative interactions is believed key to ensure that robust development emerges from a backdrop of stochastic gene expression and widespread, transient TF binding (Spitz and Furlong, 2012).

Lineage specification in early development proceeds from progenitor cells which are permissive for the initial stages of differentiation but preserved in a state of self-renewal for a number cell cycles. *Ex vivo*, these progenitor cells can be trapped indefinitely in culture as stem cells, implying that there are intrinsic mechanisms that can be exploited to block commitment while preserving differentiation potential. In the hematopoietic system, it has been suggested that self-renewal and differentiation competence is governed by the competition between lineage specific TFs in a phenomenon referred to as multi-lineage priming (Hu et al., 1997). Here, we consider how mechanisms for lineage segregation encoded in TF stoichiometry and cooperative interactions can be exploited to support embryonic stem cells’ (ESCs) self-renewal.

Naïve ESCs are a cell culture system derived from the *ex vivo* expansion of the preimplantation embryo. They are immortal, self-renewing cultures endowed with the capacity to differentiate into all lineages of the future embryo, a property known as pluripotency (Evans and Kaufman, 1981; Martin, 1981). ESCs are maintained *in vitro* by a combination of extrinsic signaling and a network of master TFs including OCT4, SOX2 and NANOG (Chambers et al., 2007; Li et al., 2007; Mitsui et al., 2003; Nichols et al., 1998). The prevailing model is that these TFs, referred to as the pluripotency network, act in a concerted feed forward loop to maintain an undifferentiated state through mutual induction and repression (Jaenisch and Young, 2008; Silva and Smith, 2008; Young, 2011). However, several of these TFs have distinct roles in pre-implantation development and later differentiation (Avilion et al., 2003; Frum et al., 2013; Mitsui et al., 2003; Nichols et al., 1998; Yamaguchi et al., 2005; Zhang et al., 2018).

While pluripotency exists indefinitely *in vitro*, *in vivo* it is a transient characteristic of the stage from which ESCs are derived. Mammalian preimplantation development begins following fertilization with a series of cleavage divisions and segregation into three distinct lineages, the Trophectoderm (TE), the Primitive Endoderm (PrE) and the Epiblast (Epi), which are the founder lineages of the placenta, the yolk sac, and the embryo proper, respectively. In mouse, these three lineages are formed as a result of two successive lineage choices: First the outer cells are specified to the TE lineage, and second the PrE and Epi are resolved from a bipotent progenitor pool of inside cells, termed the inner cell mass (ICM) (Chazaud et al., 2006; Plusa et al., 2008). The ICM segregation is driven by lineage determinant TFs, including SOX2 and NANOG in the Epi, and GATA6 in the PrE (Avilion et al., 2003; Bessonnard et al., 2014; Cai et al., 2008; Frankenberg et al., 2011; Messerschmidt and Kemler, 2010; Mitsui et al., 2003) with FGF/ERK signaling delineating the choice between Epi and PrE identity (Ambrosetti et al., 1997; Chazaud et al., 2006; Frankenberg et al., 2011; Kang et al., 2013; Yamanaka et al., 2010). During early phases of lineage specification, lineage-biased ICM cells retain the capacity to switch identity when challenged in heterochronic grafting experiments (Grabarek et al., 2012). In this paper, we define this window of plasticity as a phase of reversible lineage priming, distinct from the subsequent phase where cells commit to their chosen fate. The duration and timing of FGF/ERK signaling controls commitment and is therefore fundamental to ensuring the correct proportions of Epi and PrE cells and as such the robustness of pre-implantation development (Saiz et al., 2020; Yamanaka et al., 2010).

A number of studies in other models suggest that the interpretation of signaling duration is an emergent property of the gene regulatory network (GRN) downstream of the signal (Jutras-Dubé et al., 2018). In the context of ESCs, we have found that early ERK-mediated differentiation is reversible and depends on sustained expression of key members of the pluripotency GRN (Hamilton et al., 2019). Moreover, the ability of cells to re-establish pluripotent gene activity following a burst in ERK signaling is dependent on ESRRB (Estrogen Related Receptor Beta). Several studies have shown the importance of ESRRB in naïve pluripotency, including demonstrations that loss of ESRRB causes compromised ESC fitness, and its overexpression can substitute for NANOG function in establishing and supporting pluripotency (Festuccia et al., 2012, 2016, 2018b, 2018a). However, there is also *in vitro* evidence supporting an active role for ESRRB in PrE differentiation as well as direct upregulation of the PrE marker *Gata6* (Wamaitha et al., 2015; Uranishi et al., 2016; Herchcovici Levy et al., 2022). Does ESRRB promote differentiation at the same time as safeguarding pluripotency, and how can these two affiliations be reconciled?

In this paper, we consider whether differential stoichiometry between two TFs can underlie phenotypic shifts between self-renewal and early differentiation, providing a means to safeguard plasticity as cells progress in lineage specification. We focus on ESRRB and ask whether it solely functions as a pluripotency factor by virtue of its interaction with NANOG. We find that when both factors are expressed at physiological levels, ESRRB supports pluripotency. However, when NANOG is not present, ESRRB induces spontaneous PrE differentiation ancillary to supporting pluripotency. Based on these observations, we provide a theoretical framework for how cooperative interactions between lineage-affiliated TFs enable cells to integrate environmental cues over time as they sit at the cusp of differentiation.

## Results

### Esrrb regulates early PrE specification

While ESRRB has been implicated in PrE differentiation (Herchcovici Levy et al., 2022; Wamaitha et al., 2015) its primary described function during pre-implantation development is in naïve pluripotency (Festuccia et al., 2012, 2016, 2018b, 2021). To explore the role of ESRRB in ICM segregation, we assessed the expression of this factor in the embryonic day (E)3.5 and E4.5 blastocyst through reanalysis of previously published single cell sequencing (Nowotschin et al., 2019). To get an optimal time estimate for lineage specification, we ran the scVelo dynamical model, where the ratio between spliced and unspliced transcript is used to estimate the time derivative (RNA-velocity) of the gene expression state in an individual cell (Bergen et al., 2020; La Manno et al., 2018) (Figure 1A). This analysis provides an estimate of differentiation directionality and latent time, a parameter that captures the developmental time with precision superior to traditional pseudotime (Bergen et al., 2020) (Figures S1A-C). When assessing gene expression originating in the unspecified ICM along either latent time (Figure 1B) or pseudotime (Figure S1D), we found that *Esrrb* was expressed early in the PrE lineage, namely in the E3.5 PrE founder cells that also express the canonical markers *Gata6*, *Pdgfrα* and *Gata4* (Figures 1B and S1D), consistent with previous findings linking ESRRB to specific PrE marker genes (Uranishi et al., 2016; Wamaitha et al., 2015). As Nanog and Esrrb are known to have an essential partnership in ESCs (Festuccia et al., 2012, 2018a) and Nanog is a core PrE antagonist (Mitsui et al., 2003; Singh et al., 2007), we also looked at *Nanog* expression in the early PrE and found it was downregulated notably before *Esrrb* (Figures 1B, S1D and (Boroviak et al., 2014)). This contrasted the patterns of regulation for *Esrrb* and *Nanog* during TE and Epi differentiation, where the two factors are downregulated in parallel as lineage specific markers are initially expressed (Figures 1B and S1D). Based on this observation, we asked if ESRRB was also expressed in the absence of NANOG during PrE differentiation *in vitro*. Using defined conditions for *in vitro* PrE differentiation that closely mimics *in vivo* lineage specification (Anderson et al., 2017; Perera et al., 2022) we directed cells towards PrE and analyzed for ESRRB, NANOG and GATA6 expression. Figure 1C demonstrates the presence of a PrE founder population that expresses both GATA6 and ESRRB, but not NANOG, at day 3 of differentiation. To quantify whether this was true for all PrE founders, we generated an ESRRB::tdTomato, NANOG::EGFP double reporter cell line (Figures S2A-K). During differentiation to PrE, we observed a marked difference in the decay of ESRRB and NANOG: On day 3 of *in vitro* differentiation, we found that all cells positive for PDGFRα also expressed ESRRB but had downregulated NANOG (Figure 1D), resembling the *in vivo* expression pattern of early PrE. As these observations support the existence of a transient state in differentiation that depends on differential rates of downregulation, we carefully assessed the half-life of these fusion proteins and found that they accurately reflect the respective endogenous proteins (Figures S2L, M).

**Figure 1.**
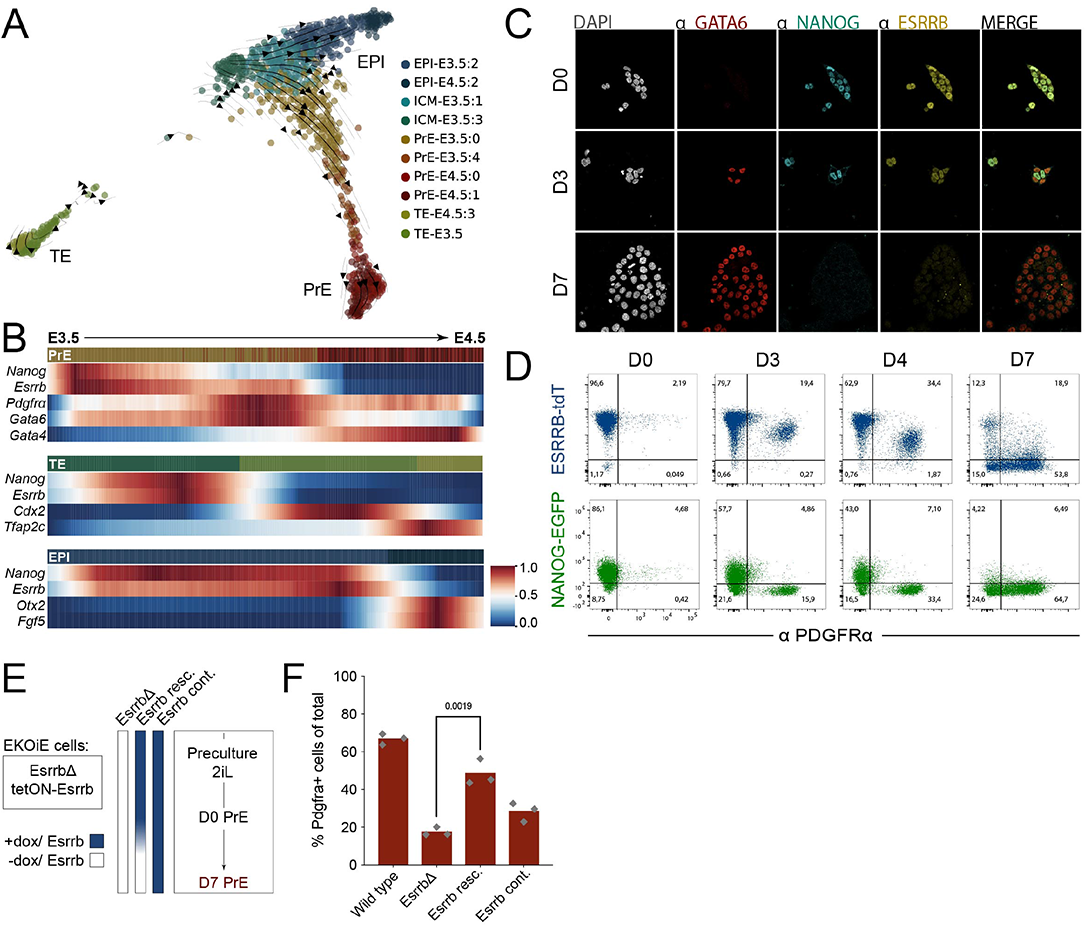
Esrrb regulates early PrE specification. **(A)** t-distributed stochastic neighbor (tSNE) embedding of complete mouse blastocyst single cell sequencing at E3.5 and E4.5 (Nowotschin et al., 2019). Clusters are based on published annotation and projected RNA-velocity was generated using scVelo (Bergen et al., 2020). **(B)** Heatmaps of scaled expression of indicated genes along scVelo defined latent time (colored bar at the top of each subplot, colors as in A) for PrE (top), TE (middle) and Epi (bottom). **(C)** Immunofluorescent imaging of the indicated markers across *in vitro* PrE differentiation (D = day, scalebar = 20μm). **(D)** Representative flow cytometry plots of ESRRB::tdT (tandemTomato) and NANOG::EGFP fusion protein expression and PDGFRα antibody staining across *in vitro* PrE differentiation, n = 3, biological clones (D = day). **(E)** Schematic of *in vitro* PrE differentiation in Esrrb conditional knock out cells, EKOiE (*Esrrb^-/-^:tetON-Esrrb*) (Festuccia et al., 2016). In the EsrrbΔ condition, *Esrrb* was not induced at any point, Esrrb rescue had *Esrrb* transgene induction until differentiation start, and Esrrb continuous had transgene induction before and throughout differentiation. **(F)** Flow cytometry analysis of the PrE marker, PDGFRα, expression at D7 of *in vitro* PrE differentiation for the indicated genotypes/ conditions (unpaired t-test was used for statistical analysis, n = 3).

The persistent expression of ESRRB in all PrE founders suggests that Esrrb is required for the early phases of *in vitro* PrE differentiation. To test this hypothesis, we performed PrE differentiation using an Esrrb knock out cell line rescued with a doxycycline (DOX) inducible *Esrrb* transgene, EKOiE (Figure 1E) (Festuccia et al., 2016). Loss of ESRRB (EsrrbΔ) significantly impaired PrE differentiation efficiency, but the transgenic expression of ESRRB before and briefly into differentiation (Esrrb resc.) significantly improved this efficiency (Figures 1F and S1E, F). This contrasts with the continuous expression of ESRRB (Esrrb cont.) which blocks PrE differentiation, as evidenced by the maintained expression of ESC marker PECAM in ∼60% of cells in these cultures (Figures 1F and S1G).

To test whether the role of ESRRB was specific to PrE differentiation or if the observed differentiation phenotype was caused by a more general impact of ESRRB on ESC fitness, we challenged the cells to neural differentiation. We observed no effect on the efficiency of neural differentiation upon Esrrb removal (Figures S3A-C), yet as with PrE differentiation, extended ESRRB expression blocked neural differentiation and supported continuous expression of ESC genes throughout differentiation (Figures S3A-C). These results indicate a context-dependent dual role for ESRRB in maintaining self-renewal and promoting PrE differentiation.

### ESRRB supports both ESC/Epi-like and PrE-like identity in the absence of NANOG

Since NANOG is known to direct the binding of key pluripotency TFs, including ESRRB, in ESCs (Heurtier et al., 2019) and is also a potent inhibitor of the commitment of ICM cells to PrE *in vivo*, we hypothesized that a PrE-specific activity of ESRRB could be regulated by NANOG. As ESRRB is robustly downregulated in NanogΔ ESCs (Festuccia et al., 2012), we could rescue it to physiological levels while uncoupled from transcriptional regulation by NANOG and hence recapitulate the ESRRB:NANOG stoichiometry observed in PrE founders. We rescued NanogΔ cells, TβC44Cre6 (Chambers et al., 2007; Navarro et al., 2012) with either the short (ES) or long (EL) isoform of Esrrb, or as controls either the short isoform of Nanog (NS) or an empty vector (EV). We used wild type (WT) E14 ESCs cultured in Serum/LIF and transfected with EV as a reference to identify stable NanogΔ cell lines that expressed the respective transgene at physiological levels (Figures 2A, B). All lines rescued with either ESRRB or NANOG regained ESC morphology. To assess the nature of steady state Serum/LIF-supported culture in cells expressing ESRRB in the absence of NANOG, we analyzed levels of the ESC marker PECAM and the PrE marker PDGFRα by flow cytometry. Figure 2C shows that *Nanog* mutants rescued by either ESRRB isoform exhibit robust upregulation of PDGFRα in a subpopulation of the culture, not observed in either control ESCs or when NANOG was reintroduced (Figure 2C). This phenotype is also reflected upon reinspection of published data in which NANOG activity is rescued by ESRRB (Festuccia et al., 2012 Figure 5).

**Figure 2.**
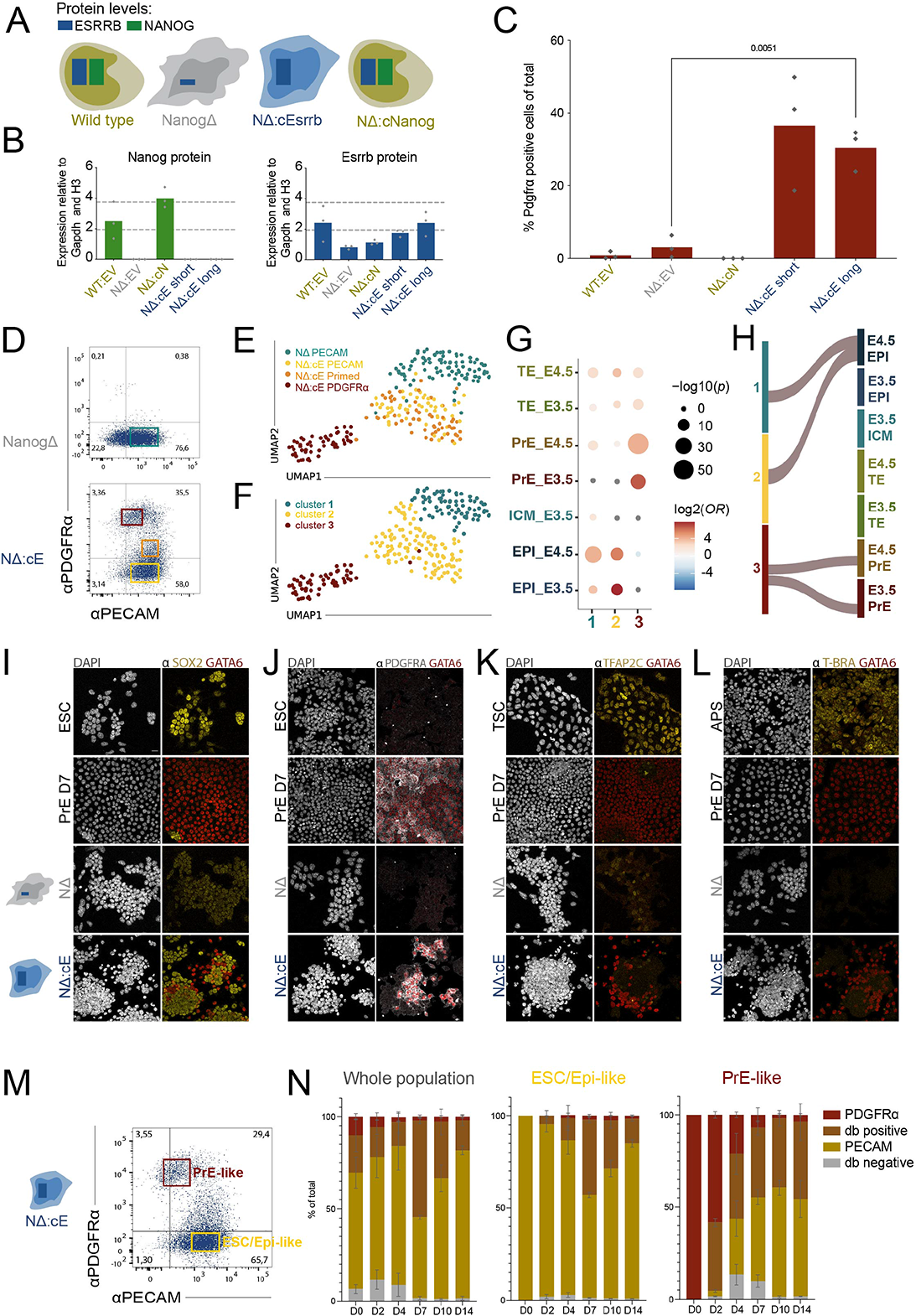
ESRRB supports both ESC/Epi-like and PrE-like identity in the absence of NANOG. **(A)** Schematic representation of the genotypes tested and their level of ESRRB (blue) and NANOG (green) protein **(B)** Quantification of NANOG and ESRRB protein levels across the indicated genotypes cultured in SL, quantified from western blotting of 3 biological clones pr genotype. WT = wild type, EV = empty vector, cN = pCAG-Nanog (short isoform cDNA), cE = pCAG-Esrrb (short and long isoform cDNA respectively). **(C)** Flow cytometry analysis of the PrE marker, PDGFRα, expression across genotypes, cultured in SL. Abbreviations as in B. (unpaired t-test was used for statistical analysis, n = 3). **(D)** Flow cytometry sorting strategy for single cell RNA sequencing of SL cultured NanogΔ and NanogΔ:cEsrrb cells stained for the PrE marker PDGFRα and the ESC marker PECAM. Cells were sorted from 3 biological replicates. **(E)** Uniform Manifold Approximation and Projection (UMAP) dimensionality reduction of single cell transcriptome of the genotypes and populations indicated. Colors correspond to the gates indicated by colors in D. **(F)** Louvain clustering of E. For cell type contribution across the clusters see Figure S3A. **(G)** GeneOverlap analysis of clusters in F against marker lists from *in vivo* single cell RNA-seq of pre-implantation blastocysts (Nowotschin et al., 2019). Grey dots indicate Odds Ratio (OR) of 0. **(H)** Analysis of cluster identity by CAT between clusters identified in F and published single cell transcriptome data of pre-implantation blastocysts. **(I-L)** Immunofluorescent staining and imaging of indicated cell types and markers. ESC = wild type ESCs in SL conditions, PrE D7 = end point of *in vitro* PrE differentiation, TSC = Trophoblast Stem Cells, APS = 48hr timepoint of anterior primitive streak differentiation. Scalebar in H = 20uM, representative for all panels. **(M)** PrE-like and ESC/Epi-like cells sorted by flow cytometry from NanogΔ:cEsrrb cells stained for the PrE marker, PDGFRα, and the ESC marker, PECAM (n = 3, biological replicates). **(N)** Sorted populations as outlined in M as well as a sorted control (whole population), analyzed across 14 days in SL culture by flow cytometry every 2^nd^ or 3^rd^ day. Cells were stained for the PrE marker, PDGFRα, and the ESC marker, PECAM.

To better understand the identity of cells expressing ESRRB in the absence of NANOG, we conducted massively parallel single cell sequencing (MARS-seq2) (Keren-Shaul et al., 2019) of specific populations in NanogΔ:Esrrb cells and the parent NanogΔ cells. We sorted PECAM^+^ cells that did not express PDGFRα from both genotypes, and from the NanogΔ:cEsrrb cells we further sorted PECAM^+^PDGFRα^low^ cells (‘primed’) as well as PDGFRα^high^ cells (Figure 2D). Dimensionality reduction places the cells on a continuum with NanogΔ PECAM^+^ cells at one end, NanogΔ:cEsrrb PECAM^+^ cells in the middle and NanogΔ:cEsrrb PDGFRα^high^ cells at the other end (Figure 2E). The NanogΔ:cEsrrb PECAM^+^ and ‘primed’ cells appear indistinguishable (Figure 2E), suggesting that the gene expression changes that enable the transition to the PDGFRα^high^ state occurred upon rescue of NanogΔ cells with ESRRB.

To infer cell identity across the populations in the dataset, we generated Louvain clustering of the single cell data (Figures 2F and S4A) and extracted marker genes for all clusters (Table S1). We compared these gene sets to marker genes of pre-implantation and post-gastrulation single cell embryo sequencing (Nowotschin et al., 2019; Pijuan-Sala et al., 2019) (Table S2) using odds ratio and p-value for significance of gene overlap between marker lists (Figures 2G and S4B; Table S3). We see increased overlap between pre-implantation Epi marker genes and the PECAM^+^ populations upon rescue of NanogΔ cells with Esrrb (Figure 2G). In addition, we observed a specific overlap between the PDGFRα^high^ cells and both E3.5 and E4.5 *in vivo* PrE (Figure 2G). To eliminate the possibility of introducing a bias by looking at only a subset of marker genes, we also used the recently developed Cluster Alignment Tool (CAT) (Rothová et al., 2022), that allows comparison of cell identities across datasets and sequencing technologies. CAT identifies relatedness of predefined clusters using an estimated average expression of all genes per cluster to calculate Euclidian distances. When comparing overall cluster identities between the clusters in our dataset and the clusters from *in vivo* single cell sequencing of the blastocyst (Nowotschin et al., 2019), CAT relates NanogΔ and NanogΔ:cEsrrb PECAM^+^ cells to Epi, and NanogΔ:cEsrrb PDGFRα^high^ cells to PrE (Figure 2H; Table S4). Taken together, the combination of marker gene overlap and global cluster alignment suggests that ESRRB in the absence of NANOG promotes a PrE-like identity ancillary to supporting a pluripotent ESC/Epi-like identity.

To confirm the putative PrE identity of this sup-population, we stained NanogΔ:cEsrrb cells for canonical marker proteins of both pre- and post-implantation cell identities. We also included relevant *in vitro* differentiated cell types or stable cell lines with expression of the proteins of interest as positive controls. In NanogΔ:cEsrrb cells, GATA6 expression was evident both within and around colonies of SOX2 positive cells, but the expression of the two markers were generally mutually exclusive with no cells co-expressing high levels of both markers (Figure 2I). The GATA6 positive population that arises in the NanogΔ:cEsrrb cells also express PDGFRα, but do not express the TE or mesoderm markers TFAP2C, CDX2, T or TBX6 (Figures 2J-L and S4C-E). We also stained for neural markers TUJ1 and SOX1, as well as EpiSC markers NCAD, OCT6 and OTX2, but we observed no specific upregulation of these markers in NanogΔ:cEsrrb cells compared to WT ESCs (Figures S4F-I). We conclude that the PDGFRα^high^/ GATA6^+^ population in NanogΔ:cEsrrb cultures bears the greatest resemblance to pre-implantation PrE and that ESRRB in the absence of NANOG supports a bistable culture of bona fide ESC/Epi-like (PECAM^+^) and PrE-like (PDGFRα^high^) cells.

The stable coexistence of both Epi and PrE-like cells *in vitro* made us wonder whether constitutive expression of ESRRB in the absence of NANOG not only allowed spontaneous differentiation to PrE-like cells, but also the reverse. To address this question, we isolated cells of both identities from NanogΔ:cEsrrb cultures and plated the sorted populations back into Serum/LIF (Figures 2M, N). Already after 2 days, the PrE-like population was arising from the ESC/Epi-like cells, and strikingly, also vice versa (Figure 2N). After a week, both sorted cell types had regenerated the respective lost population and was comparable to the control of sorting the whole population (Figure 2N). In contrast, PrE cells sorted at the end of *in vitro* differentiation could not revert to the PECAM-positive ESC state when plated back into Serum/LIF culture (Figure S4J). We conclude that in the absence of NANOG, ESRRB expressed at WT endogenous levels allows cells to both take on PrE-like identity and revert back to ESC/Epi-like identity and as such dynamically interconvert, resembling the *in vivo* early Epi and PrE during the reversible lineage priming of the ICM.

### Titration of NANOG to ESRRB controls cell type specification

Next, we wished to understand and study the molecular relationship between NANOG and ESRRB, and how the relative levels of these two factors regulate cell fate. We built an *in vitro* model where we could recapitulate the NANOG and ESRRB stoichiometry of both PrE and Epi founders by constitutive expression of *Esrrb* and inducible expression of *Nanog* in the context of *Nanog* mutants. We first introduced a constitutively expressed transgene encoding the short *Esrrb* isoform linked to a self-cleaving T2A peptide (Ryan et al., 1991), followed by the coding sequence for the reverse tet activator (rtTA) (Zhou et al., 2006) into *Nanog* mutants (Figure 3A). We again selected stable lines that expressed ESRRB at similar levels to that observed in WT E14 (Figure 3B). Following this we introduced a second transgene encoding the short *Nanog* isoform under dual regulation by a DOX inducible promoter (tetON) and the destabilization tag FKBP, which can be stabilized through binding of the small molecule Shield (Shld) (Banaszynski et al., 2006; Zhou et al., 2006) (Figure 3A). The resulting cell line expressed ESRRB at endogenous levels and an inducible NANOG that could be titrated based on the addition of DOX and Shld (Figures 3A-C). The cell line is referred to as NΔ:cE:iN, and the condition of ESRRB in the absence (-DOX and Shld) or presence (+DOX and Shld) of NANOG as NΔ:E and NΔ:E:N, respectively (Figure 3C).

**Figure 3.**
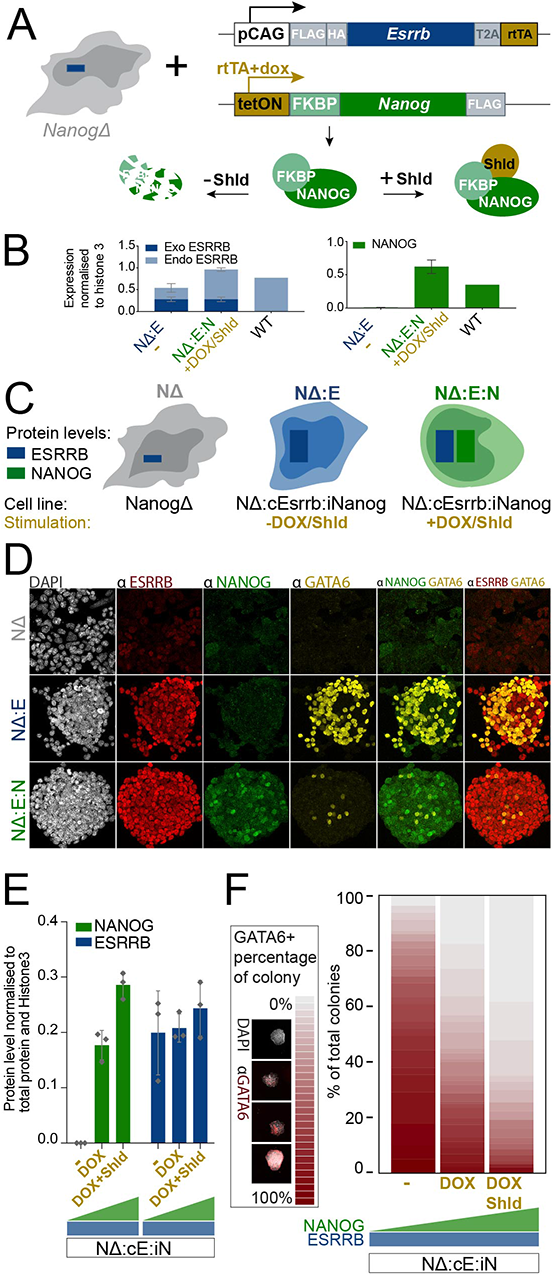
Titration of NANOG to ESRRB controls cell type specification. **(A)** Schematic representation of the ESRRB-NANOG titration cell line: *NanogΔ* cells with constitutive expression of *Esrrb* and with DOX inducible expression of *Nanog* fused to the destabilizing domain, FKBP, that targets the protein for degradation in the absence of Shld. **(B)** NANOG and ESRRB protein levels in the titration cell line schematized in A and in a WT control. Levels were quantified from western blotting of 3 biological replicates. WT = wild type, NΔ:cE = NanogΔ:cEsrrb and NΔ:cE:N = NanogΔ:cEsrrb:Nanog refers to the titration line minus and plus induction of NANOG respectively. Exo = exogenous/ transgene protein, Endo = endogenous protein. **(C)** Schematic representation of ESRRB (blue) and NANOG (green) levels across the indicated conditions (top) and genotypes (bottom). **(D)** Immunofluorescent imaging analysis of the indicated markers for the genotypes/ conditions schematised in C. **(E-F)** NΔ:cE:iN cells were cultured 24hrs without NANOG induction (-), with intermediate induction of NANOG (DOX) or with full induction of NANOG (DOX+Shld). Cells were then sorted by flow cytometry for PECAM+ PDGFRα-ESC/Epi-like cells and plated as single cells for colony expansion in the respective NANOG induction conditions they had prior to sorting. At the time of sorting, protein levels of ESRRB and NANOG across the conditions was quantified by western blotting (E). After expansion the resulting colonies were analysis by immunofluorescent imaging for GATA6 expression (F). Stacked histogram across NANOG induction conditions where GATA6+ percentage of the colony is reflected in grey to red coloring, divided into 30 bins (n = 3, biological replicates and total number of colonies analyzed pr condition was 80, 95 and 132).

Similar to our previous system, where the proteins were expressed without tags, in the context of this new cell line, NΔ:cE:iN, cells spontaneously generated a PrE-like subpopulation when ESRRB was expressed in the absence of NANOG. However, when NANOG was induced and stabilized by both DOX and Shld, this population was almost completely eliminated, alluding to cooperative interactions between the two factors (Figure 3D). To assess how the ratio of NANOG to ESRRB influenced the choice of single cells to differentiate, we performed a single cell colony expansion assay. We sorted single ESC/Epi-like cells from the NΔ:cE:iN line and plated them either devoid of NANOG, with intermediate levels of NANOG (only DOX) or with maximal NANOG (DOX and Shld) in the context of regular Serum/LIF culture conditions (Figure 3E). Each treatment was initiated 24 hours before sorting and continued after sorting during colony expansion. The resulting colonies were then fixed, stained for GATA6, imaged and analyzed using an automated pipeline for quantification of the GATA6 positive fraction of the colonies. The assay showed that the upregulation of the PrE marker GATA6 is inversely proportional to the level of NANOG (Figure 3F). Without NANOG, ∼70% of colonies were >50% GATA6 positive, with intermediate or fully induced NANOG this was true for ∼30% and ∼10% of colonies, respectively (Figure 3F). Taken together, these observations support the idea that the ratio of ESRRB to NANOG determines the likelihood that a cell will differentiate to the PrE-like state or maintain itself in the ESC/Epi-like pluripotent state.

### The transcriptional basis of ESRRB-dependent PrE priming

To understand why the ratio of NANOG to ESRRB influences cell fate choice, we focused on how ESRRB influenced low-level transcription and chromatin accessibility in the ESC/Epi-like cells with potential to form PrE-like cells. We therefore sorted this population from both non-induced and induced NΔ:cE:iN cells (NΔ:E and NΔ:E:N, respectively) as well as from NanogΔ parent cells (NΔ), and we conducted bulk RNA and Assay for Transposase-Accessible Chromatin (ATAC) sequencing (Figure 4A). Principle component analysis of the resulting transcriptomic data primarily separated the NΔ cells from NΔ:E and NΔ:E:N (PC1, ∼75% of all variance) (Figure 4B). Differential gene expression analysis identified 1398, 346 and 474 uniquely upregulated genes for NΔ, NΔ:E and NΔ:E:N, respectively (Table S5; Log2FC > 1 and adjusted p-value < 0.01). Using these gene sets we explored how they compared to the gene expression observed *in vivo* in pre-implantation development (Figures 4C and S5A, B; Table S6). We found that both NΔ:E and NΔ:E:N upregulate E3.5 Epi markers. In addition, NΔ:E uniquely expressed E3.5 PrE markers, despite their undifferentiated ESC phenotype. This observation indicates that even in the ESC/Epi-like population of NΔ:E cells, the expression of ESRRB in the absence of NANOG is priming cells for PrE-like differentiation.

**Figure 4.**
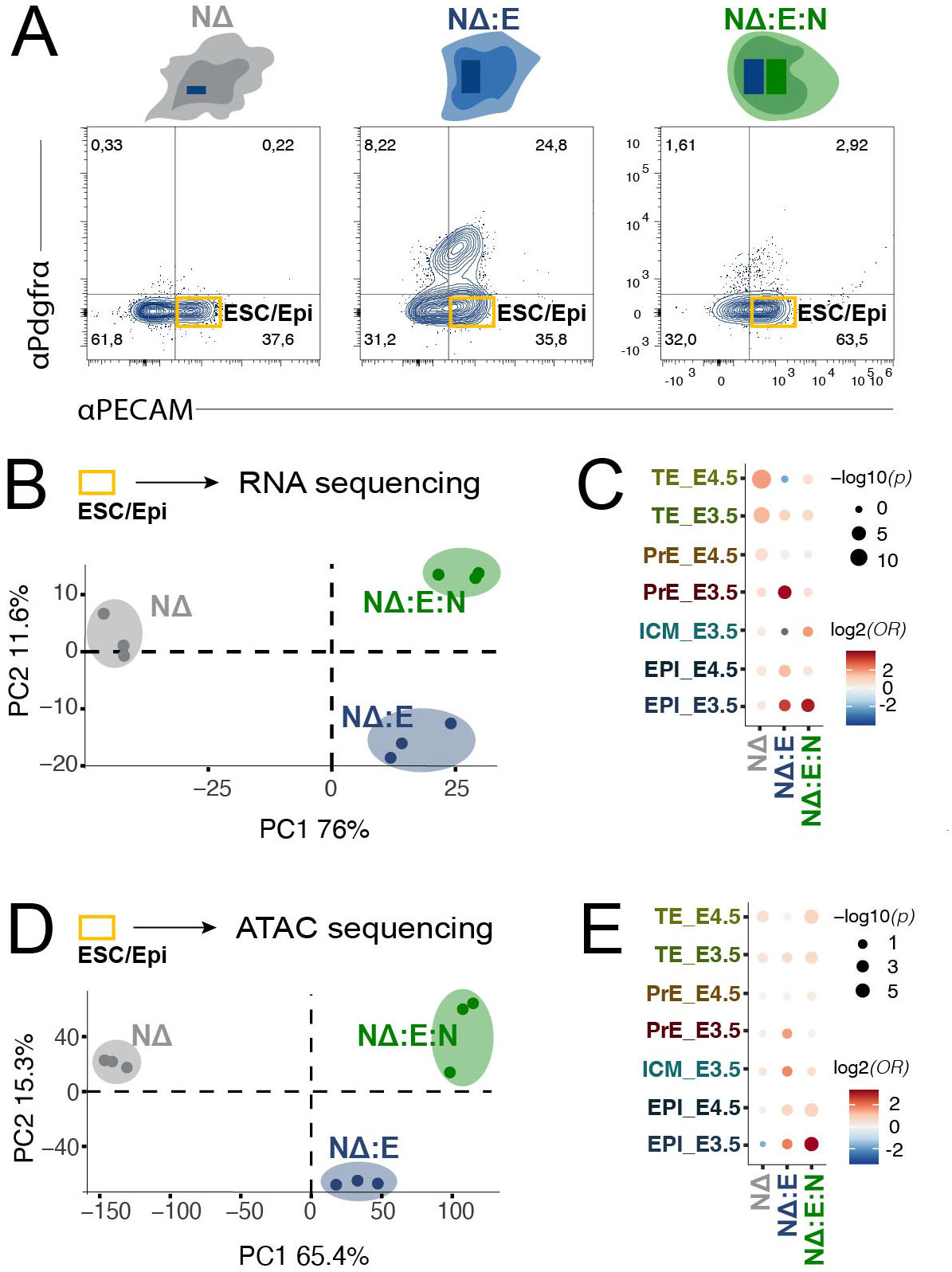
The transcriptional basis of ESRRB-dependent PrE priming. **(A)** Representative flow cytometry plots of sorting strategy prior to ATAC- and RNA-sequencing of NanogΔ cells and NanogΔ:cEsrrb:iNanog cells without and with Nanog, NΔ:E and NΔ:E:N respectively. Cells were stained for the PrE marker, PDGFRα, and the ESC marker, PECAM, ESC/Epi-like cells, and sorted based on the yellow gates on the plots (n=3, biological replicates). **(B)** Principal Component Analysis (PCA) of the transcriptome variance across cells sorted in A. **(C)** GeneOverlap analysis of condition specific differentially expressed genes (L2FC > 1 and adjusted p-value < 0.01) against marker lists from *in vivo* single cell RNA-seq of pre-implantation blastocysts (Nowotschin et al., 2019). Grey dots indicate OR of 0. **(D)** PCA of chromatin accessibility variance across cells sorted in A. **(E)** GeneOverlap analysis of condition specific differentially open peaks (L2FC > 1 and adjusted p-value < 0.01) annotated to nearest gene against marker lists from *in vivo* single cell RNA seq of pre-implantation blastocysts (Nowotschin et al., 2019).

The observation that these cells are drifting toward PrE is also reflected in their chromatin, which exhibits an overall distribution of variance similar to the transcriptome (Figure 4D). From 157498 consensus peaks, we extracted 11893, 2172 and 8879 peaks uniquely upregulated in NΔ, NΔ:E and NΔ:E:N respectively, (Figures S5C, D; Table S7; Log2FC > 1 and adjusted p-value < 0.01). We annotated the peaks to their nearest genes and compared these genotype-specific gene sets to pre-implantation development (Figure 4E; Tables S8, 9). Genomic regions with enriched chromatin accessibility in NΔ:E ESC/Epi-like cells were primarily close to marker genes of E3.5 PrE, ICM and Epi, whereas regions enriched in the NΔ:E:N condition were in proximity to genes that define E3.5 Epi (Figure 4E). Given that our clonal data shows a propensity to make PrE in ∼90% of NΔ:E ESC/Epi-like cells (Figure 3F), we conclude that these cells represent a single uniform population with a chromatin landscape permissive for both self-renewal and differentiation.

We conducted motif analysis of the identified enriched ATAC regions for each of the 3 conditions and found the Esrrb motif in both NΔ:E and NΔ:E:N, where it was present in 24 and 26% of the peaks, respectively (Figure S5E; Table S10). In addition, we observed an enrichment of Oct4-Sox2-Tcf-Nanog and KLF motifs coupled to the reintroduction of NANOG. This observation suggests that cooperative interaction between ESRRB and NANOG might be causing the ESC/Epi-like promoting activity of ESRRB in this context.

### Esrrb regulates PrE genes by direct DNA binding

To understand the specific role of ESRRB activity in the observed changes on chromatin and transcription, we assessed ESRRB binding in the presence or absence of NANOG by ChIP-sequencing. For the vast majority of binding sites, ESRRB occupancy was not affected by the presence or absence of NANOG, however a significant shift in binding was observed for 5,3% of the binding sites (out of 86.202 consensus peaks we found 1664 and 2892 peaks enriched in NΔ:E and NΔ:E:N respectively, using a cutoff of absolute Log2FC > 1 and adjusted p-value < 0.05) (Figures 5A and S6A; Table S11). Peaks enriched for ESRRB binding both in the presence or absence of NANOG had high prevalence of the Esrrb motif, but the Oct4-Sox2-Tcf-Nanog motif was uniquely prevalent in the peaks enriched for ESRRB binding in the presence of NANOG (Figure 5B; Table S12), suggesting that ESRRB binding at these sites was stabilized by cooperative binding with NANOG. In contrast, ESRRB binding in the absence of NANOG only resulted in a small overrepresentation of Gata motifs (Figure 5B).

**Figure 5.**
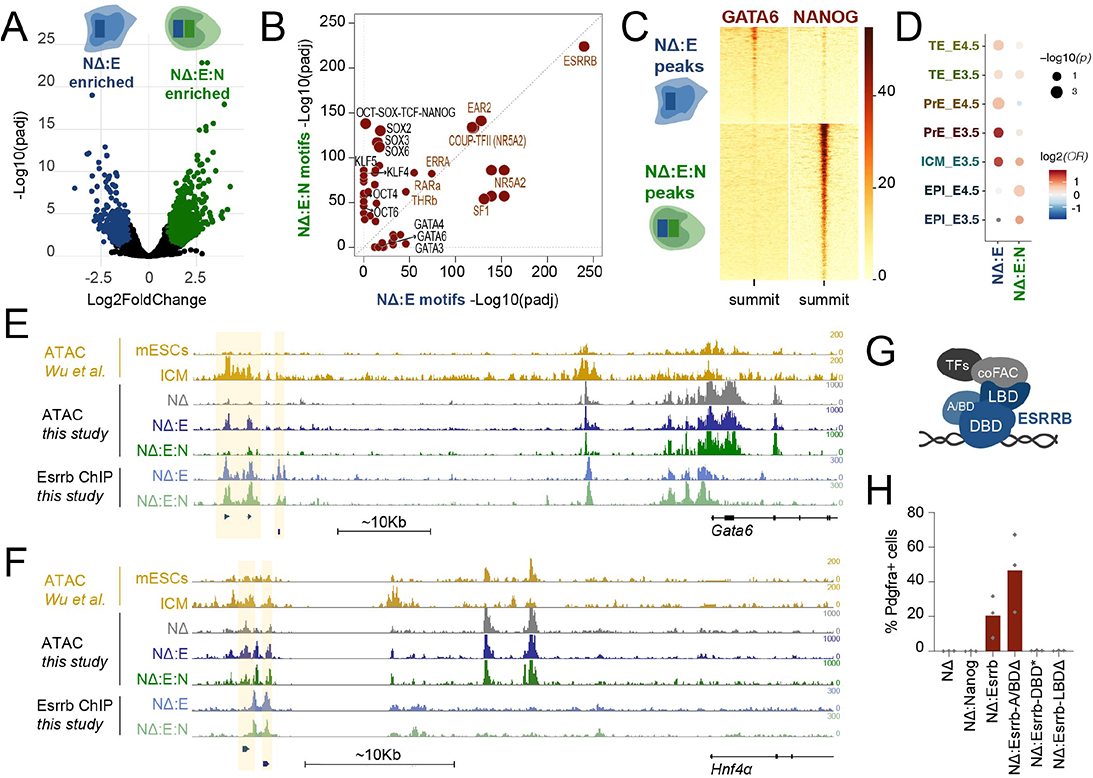
Esrrb regulates PrE genes by direct DNA binding. **(A)** Volcano plot of ESRRB Chromatin Immunoprecipitation (ChIP) sequencing peaks and enrichment in NanogΔ:cE:iN cells without and with Nanog induction, NΔ:E and NΔ:E:N respectively. Peaks differentially enriched between conditions with absolute L2FC > 1 and adjusted p-value < 0.05 are colored (n=3, biological replicates). **(B)** Scatter plot of motif enrichment in peaks with significantly more ESRRB binding in the absence of NANOG (NΔ:cE) versus in the presence of NANOG (NΔ:cE:N) (thresholds as in A). Nuclear receptors are labelled with red text. **(C)** Binding profiles of GATA6 (Wamaitha et al., 2015) and NANOG (Marson et al., 2008) at either peak class defined in A. **(D)** GeneOverlap analysis of condition specific differentially bound ESRRB peaks (thresholds as in A) annotated to nearest gene against marker lists from *in vivo* single cell RNA-seq of pre-implantation blastocysts (Nowotschin et al., 2019). Grey dots indicate OR of 0. **(E-F)** RPKM normalized bigwig tracks upstream the *Gata6* (**E**) and *Hnf4α* (**F**). Tracks are ATAC sequencing in SL cultured ESCs and *in* vivo ICM (Wu et al., 2016) in yellow. ATAC sequencing from the ESC/Epi-like population of NanogΔ, NanogΔ:cEsrrb:iNanog without and with Nanog induction. ESRRB ChIP-seq in NanogΔ:cEsrrb:iNanog without and with Nanog induction. Regions of interest are highlighted and peaks significantly enriched specifically in the NΔ:E condition in the ATAC (top) and ChIP (bottom) are depicted below the tracks. **(G)** Schematic representation of ESRRB (blue) on DNA, with the DBD binding DNA and the LBD recruiting co-factors etc. A/BD = N-terminal transactivation domain, DBD = DNA Binding Domain, LBD = Ligand Binding Domain. **(H)** *NanogΔ* cells were rescued with *Nanog*, *Esrrb*, *Esrrb* with point mutations in the DBD that prohibit DNA binding or truncated forms of *Esrrb* lacking either the A/BD or LBD. Flow cytometry analysis of the PrE marker, PDGFRα, expression across genotypes (n = 3, biological clones cultured in SL).

To investigate ESRRB co-binding with NANOG and the PrE factor GATA6, we compared our data with previously published ChIP-seq for NANOG and GATA6 (Marson et al., 2008; Wamaitha et al., 2015). Both datasets were from mouse ESCs cultured in Serum/LIF, and in the case of GATA6, also induced with GATA6 expression for 36hrs, which promotes PrE-like differentiation (Wamaitha et al., 2015). The enriched ESRRB binding sites in the presence of NANOG overlapped with NANOG binding, but not GATA6. In contrast, ESRRB binding in the absence of NANOG overlapped with GATA6, suggesting that a cooperative binding with NANOG stimulates the disassociation of ESRRB from PrE affiliated genes in favor of its association with pluripotency genes (Figure 5C).

To understand the impact of these ESRRB bound regions on gene regulation and ultimately cell identity, we annotated the peaks to nearest ESRRB or NANOG upregulated gene (i.e., excluding genes uniquely upregulated in the *NanogΔ* bulk transcriptome, Table S13). We then used gene overlap analysis to uncover the prevalence of lineage associated genes. We found that the peaks enriched for ESRRB binding in the absence of NANOG were near E3.5 PrE and ICM genes, whereas peaks enriched for ESRRB binding in the presence of NANOG were near Epi and ICM marker genes (Figure 5D; Table S14). Examples of ESRRB binding and opening the chromatin landscape at PrE associated loci in the absence of NANOG are shown for the *Gata6* and *Hnf4α* loci (Figures 5E, F). In these loci we identified regulatory elements upstream of the transcription start site that were accessible in *in vivo* ICM where *Gata6* and *Hnf4α* are expressed, but closed in ESCs, where the two genes are not expressed (Wu et al., 2016). These regions were specifically open in NΔ:E ESC/Epi-like cells and both had an adjacent Esrrb binding site, where ESRRB binding was enriched in the absence of NANOG (Figures 5E,F).

Altogether, the analysis of ESRRB binding in the presence and absence of NANOG suggests a model where ESRRB, in a context dependent manner, promotes two distinct cell identities through sequence-specific DNA binding and regulation of chromatin accessibility, which ultimately triggers changes in gene expression. To test this model, we rescued *Nanog* mutants with a panel of ESRRB mutants and assessed their ability to induce a PrE-like subpopulation in the absence of NANOG. ESRRB consists of an N terminal transactivation domain (A/BD), followed by a DNA binding domain (DBD) and a ligand binding domain (LDB). While both DNA binding and co-factor recruitment via the LBD are essential for ESRRB function in ESC self-renewal and pluripotency, the transactivation domain is dispensable (Festuccia et al., 2018b; Gearhart et al., 2003; Percharde et al., 2012; Uranishi et al., 2013). To engineer an ESRRB mutant that cannot bind DNA, the amino acids responsible for DNA binding (Glu142, Lys145 and Lys149) were mutated to glycines (DBD*); we also generated truncations that remove either the LBD (LBDΔ, removal of aa 235-457) or the A/BD (A/BDΔ, removal of aa 1-100). We introduced these mutants into *NanogΔ* cells following the strategy applied in Figures 2A and B (Figures S6B, C) and assayed the expression of PDGFRα protein by flow cytometry (Figures 5G and S6D). Similar to its activity in pluripotency, we found that functional DNA binding and co-factor recruitment via the LBD were required for ESRRB to induce PrE-like cells, while the N-terminal transactivation domain was dispensable.

### The bipartite function of ESRRB provides a molecular understanding of PrE/Epi founder plasticity

To distil the potential implication of a bipartite ESRRB activity on lineage specification, we used mathematical modeling. We focused on testing the hypothesis derived from our manipulation of ESC states, that ESRRB could be supporting an undecided, primed but not committed state, as a consequence of its ability to regulate both PrE and Epi/pluripotent identity. In embryonic development, pausing in such a state may allow cells to integrate fluctuating signals over time (e.g. FGF) and may be necessary for the embryo to adapt to alterations in its cellular composition (Grabarek et al., 2012; Yamanaka et al., 2010).

Mathematical models developed for the understanding of PrE-EPI segregation all converge on the importance of mutual antagonism between GATA6 and NANOG, coupled with paracrine FGF signaling, for the robust PrE/Epi ratio (Bessonnard et al., 2014; Chickarmane et al., 2006; De Mot et al., 2016; Nissen et al., 2017; Saiz et al., 2020; Schro ter et al., 2015; Tosenberger et al., 2017). A classical approach to model the segregation of two lineages is to use ordinary differential equations (ODEs, generalized equation is shown in Figure S7A) (Gardner et al., 2000; Alon, 2007; Huang et al., 2007). In the case of PrE-Epi segregation, the ODEs formalize the GATA6-NANOG-FGF regulatory network, and the dynamics converge on at least two stable cell identities (stable fix points), namely Epi (NANOG^high^-GATA6^low^) and PrE (NANOG^low^-GATA6^high^). There are, however, different *in silico* interpretations of the bipotent ICM state (GATA6-NANOG co-expressed): it is thought of either as a transient state that blastomeres progress through while going from NANOG^low^-GATA6^low^ to committed PrE and Epi (Saiz et al., 2020), or as an additional stable fix point in a tristable dynamic system (Bessonnard et al., 2014; De Mot et al., 2016; Tosenberger et al., 2017).

To understanding how a bipartite ESRRB activity influences a simple mutually antagonistic network, we compared the dynamics of a distilled, bistable GATA6-NANOG-FGF (*GNF*) network with a network also including ESRRB (*GNFE*) (Figures 6A, B). Consistent with the potential dual function of ESRRB, we constructed the GNFE network to include stimulation of GATA6 and NANOG by ESRRB, in addition to representing the capacity of NANOG to alter ESRRB activity. We also incorporated direct inhibition of ESRRB by FGF, although weaker than the inhibitory effect of FGF on NANOG, reflecting the differential half-life of these factors in response to FGF/ERK (Hamilton et al., 2019) and the persistent expression of ESRRB in PrE founders *in vivo* (Figure 1B). Specifically, we wanted to assess whether ESRRB would slow down the process of lineage specification and thus support reversible lineage priming before commitment.

**Figure 6.**
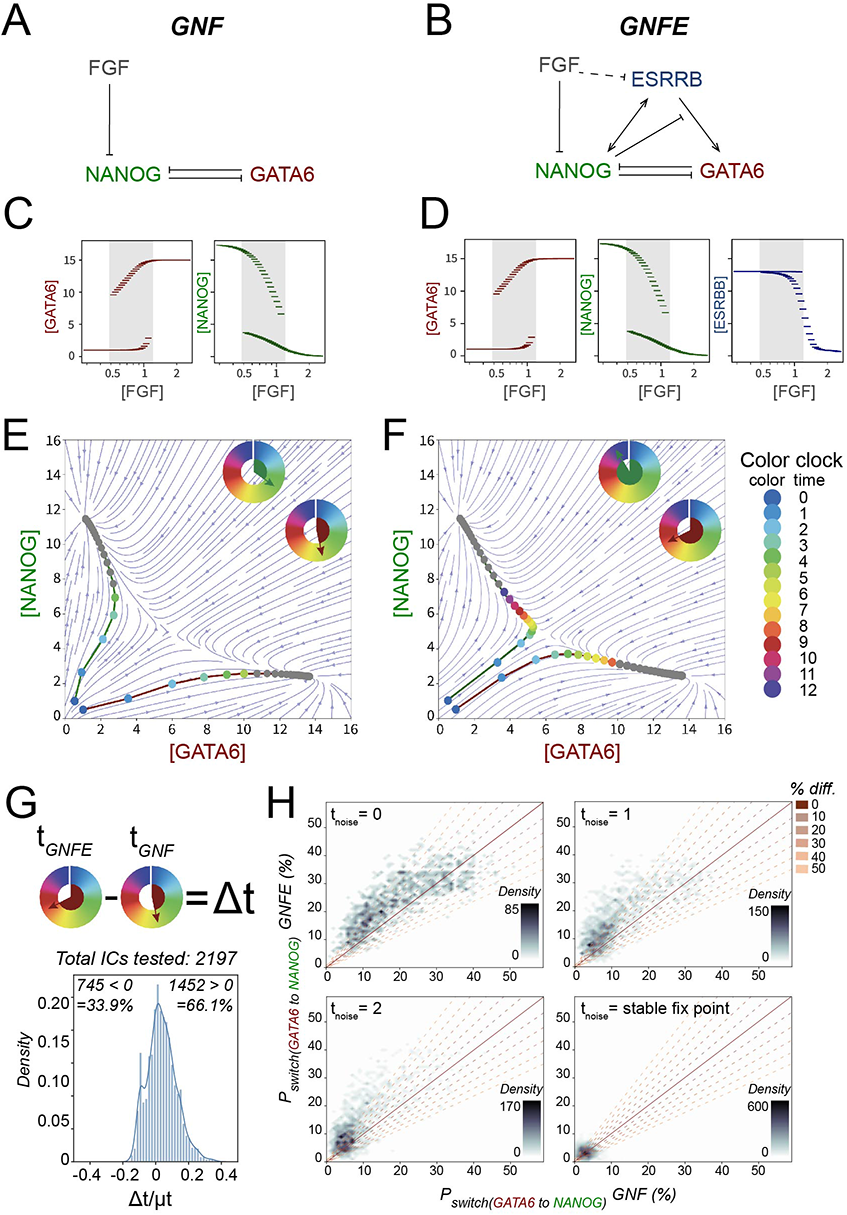
The bipartite function of ESRRB provides a molecular understanding of PrE/Epi founder plasticity. **(A-B)** Graphical representation of the GRNs explored: The simple GNF (A) with GATA6, NANOG and FGF and the more complex GNFE (B) with ESRRB added to the network. Both networks and the interactions represented are based on experimental data either presented in this study or previously published (Chazaud et al., 2006; Frankenberg et al., 2011; Fujikura et al., 2002; Hamilton et al., 2019; Nichols et al., 2009; Shimosato et al., 2007; Singh et al., 2007; Yamanaka et al., 2010). **(C-D)** Bifurcation diagrams for GNF (C) and GNFE (D) as a function of the parameter FGF, representing the FGF concentration (log-scale). Stable fix point values for GATA6, NANOG and ESRRB are marked in red, green and blue respectively, and the range of FGF concentrations supporting bistability is shaded (concurrent NANOG^high^-GATA6^low^ and GATA6^high^-NANOG^low^). **(E-F)** GATA6-NANOG phase space for FGF = 0.85, showing the vector field (blue) and the time evolutions (labelled in rainbow colors: blue through green, yellow, red to purple) from 2 initial conditions, [GATA6, NANOG] = [1,0.5] or [0.5,1]. E and F represents GNF and GNFE respectively. Dots are colored until reaching a fixed threshold (NANOG > 7.5 or GATA6 > 10). The time spent between two neighboring dots on a path is the same, and as such time can be abstractly measured in the color of the dots. The color clocks thus represent the time before reaching the threshold for NANOG (green) or GATA6 (red). Legends shows the representation of time by color. **(G)** Top: cartoon depicting the calculation of Δt as the time taken to reach the stable fix point for GNEF subtracted the time for GNF (t_GNFE_ - t_GNF_) for 2197 initial conditions in the GATA6^low^-NANOG^low^ region. Bottom: Distribution of Δt normalized to the respective average t (μt) as calculated for each initial condition. Number of initial conditions with Δt > 0 and Δt < 0 are indicated. **(H)** Scatter plots of the probability of switching fix point destination from GATA6^high^ to NANOG^high^ without (GNF) versus with ESRRB (GNFE). Final destination for each of the 2197 initial conditions was defined using the deterministic system (Figure S7B, C). Noise was introduced at 4 distinct timepoints (indicated by t_noise_) during segregation. Plots on the probability of switching from NANOG^high^ to GATA^high^ stable fix point destination is in Figure S7D.

Consistent with experimental observations (Chazaud et al., 2006; Kang et al., 2013; Nichols et al., 2009; Yamanaka et al., 2010), both networks respond to increasing FGF concentrations by transitioning from supporting Epi-only monostability, through supporting Epi-PrE bistability, to supporting PrE-only monostability (Figures 6C, D, bistable range shaded). This suggests that ESRRB does not affect the stable fix points of the system.

*In vivo* GATA6 and NANOG are initially expressed at low levels in the morula and then increase in parallel until the early blastocyst stage, where expression of NANOG and GATA6 in ICM cells becomes mutually exclusive (Chazaud et al., 2006; Plusa et al., 2008). Since we were interested in the effect of ESRRB on the lineage trajectory of a cell during blastomere specification, we compared how cells transition to the stable states of Epi or PrE in the presence or absence of ESRRB. ODE models can be displayed as a dynamic landscape (phase space) of directionality vectors that represent how any cell would transition to either of the stable fix points (Figures 6E, F). The panels show that with ESRRB the landscape changes in the GATA6^low^-NANOG^low^ region, where the trajectories with ESRRB are parallel and targeted towards the GATA6^medium^-NANOG^medium^ region, contrasting the trajectories without ESRRB which are targeted along the axes (Figures 6E, F). Hence, without ESRRB in the network, a small shift in either NANOG or GATA6 expression results in a direct path towards either stable fix point (Figure 6E, highlighted paths). However, when ESRRB is included in the GRN, lineage segregation follows a slower and more gradual path through a GATA6^medium^-NANOG^medium^ transient state before resolving into the stable cell identities (Figure 6F, highlighted paths). To quantify this globally, we ran simulations of paths taken from >2000 initial conditions in the GATA6^low^-NANOG^low^ region and measured the time it took to reach a stable fix point in either network (Figure 6G). For two thirds of the initial conditions, it took longer to reach steady state when ESRRB was present (Figure 6G), suggesting that the bipartite nature of ESRRB activity promotes gradual cell fate choice.

The implication of a gradual lineage specification event is that it could manifest as an increased ability to change fate away from the initial bias (plasticity) in response to an altered environment. To test this hypothesis, we developed stochastic versions of the two models using Gillespie’s algorithm (Gillespie, 1976), and asked whether the networks differed in their ability to adapt to changes in the local environment (represented by noise) while undergoing lineage segregation. We found that inclusion of ESRRB in the network indeed increases the probability of a cell altering fate when exposed to noise throughout lineage segregation (Figures 6H and S7D, t_noise_ = 0). Experimentally, blastomere potential becomes continuously restricted in PrE and Epi progenitors between E3.5 and 4.5 in heterochronic grafting experiments, such that they are potent to make all cells of the blastocyst at E3.5, but restricted to their respective lineage at E4.5 (Grabarek et al., 2012). We simulated this experiment by introducing noise at consecutive intermediate timepoints during lineage segregation (Figures 6H and S7D). In line with grafting experiments, the model with ESRRB showed enhanced lineage plasticity at early timepoints, yet the difference decreased with time (t_noise_ = 0,1,2). Finally, when the Epi or PrE stable fix point was reached, the two systems exhibited equivalent behaviors (Figures 6H and S7D). We conclude that *in silico* modeling supports the notion that bipartite activity of TFs, such as that described for ESRRB, could safeguard plasticity during lineage segregation, ensuring a responsive transition state, without impeding commitment.

## Discussion

In this paper we have shown that TF stoichiometry is key to expansion of ESCs and that this could originate from the need of an embryo to adapt to environmental challenges. We have focused on the pluripotency TF ESRRB and shown that it has an ancillary affiliation with the PrE lineage that is modified through interactions with NANOG to support pluripotency and self-renewal. Cooperativity between these two TFs, together with their differential response to FGF/ERK and the instructive activity of ESRRB in the early PrE, constitute a molecular network that may underpin reversible lineage priming. As such, our work suggests that *ex vivo* self-renewal exploits the developmental GRNs by stabilizing otherwise transient TF ratios.

While ESRRB has recently been defined as a pluripotency factor, it was initially identified based on its role in the TE. Collectively, a large body of work on Esrrb function across tissue types has identified a range of proliferation phenotypes. *EsrrbΔ* mice are embryonic lethal at E10.5 due to placental abnormalities and growth failure as loss of Esrrb causes extensive formation of secondary trophoblast giant cells at the expense of proliferation in the ectoplacental cone (Luo et al., 1997). In trophoblast stem cells (TSCs), ESRRB acts downstream of FGF/ERK signaling, with known partners from ESCs, as well as TSC specific interactors to block differentiation and support self-renewal (Latos et al., 2015; Tremblay et al., 2001). The TE phenotype can be rescued in WT tetraploid compensation to produce *EsrrbΔ* embryos that are viable and fertile, although these mice show behavioral phenotypes and have reduced numbers of PGCs as a result of impaired proliferation (Mitsunaga et al., 2004). An early phenotype in pre-implantation development could be masked by maternal Esrrb or a redundancy with Nr5a2 (Festuccia et al., 2021). The broad expression pattern of Esrrb in the mouse (Festuccia et al., 2018b; Petryszak et al., 2016), together with reoccurring proliferation phenotypes in different cell types, suggests a general role for Esrrb in transcriptional regulation that can be exploited in a context dependent manner to promote both self-renewal and appropriate lineage specific regulation.

This raises a general question of how altering the context changes TF activity. Here we described ESRRB activity in two contexts – with and without NANOG. We observed a strong correlation between ESRRB and NANOG binding in the context of pluripotent/Epi identity, suggesting a cooperative relationship between these TFs and a mechanism that could explain context dependent ESRRB activity. Although less pronounced, we observed similar binding alongside GATA6 in the PrE context, consistent with the cooperative interaction between GATA4 and ESRRG that was recently described in human cardiomyocytes (Sakamoto et al., 2022). Relative to the context dependent distinct modes of activity for Esrrb in ESCs and TSCs, we present a dynamic system where the presence or absence of Esrrb and Nanog is sufficient to cause distinct shifts in chromatin accessibility, transcription and ultimately cell identity.

Cooperativity between two TFs with different, but overlapping, expression patterns would represent an ideal mechanism for context dependent TF activity, that could support plasticity in the early stages of differentiation. Using ODE based modeling, we find that the presence of ESRRB, as a factor stimulating both sides of the network, enables a more gradual lineage segregation and increased responsiveness to alterations in the local environment. This is due to the transient support of a GATA6-NANOG co-expressed state, corresponding to the expression profile in the early ICM, a state that is not supported in the simple, mutually antagonistic network. However, by coupling of the simple NANOG and GATA6 network with paracrine FGF signaling, Epi-PrE segregation can be shown to proceed via tristable dynamics that includes an ICM state of NANOG-GATA6 co-expression (Bessonnard et al., 2014; De Mot et al., 2016). Alternatively, it has been suggested that the ICM state in the bistable systems could be transiently supported by an undescribed orthogonal process capable of countering the antagonistic interactions between GATA6 and NANOG and allowing blastomeres to progress through an interim state of GATA6-NANOG co-expression while going from NANOG^low^-GATA6^low^ to committed PrE and Epi (Simon et al., 2018). We have shown that bipartite TF activity affiliated with both competing lineages during segregation can support such interim “tristability” in a bistable system. Our stochastic simulations of grafting experiments (Grabarek et al., 2012) suggest that the GRN with ESRRB provides a molecular mechanism for reversible lineage priming downstream FGF/ERK signaling; namely differential degradation kinetics and bipartite TF activity bridging between two segregating lineages.

The notion that lineage segregation is governed by mutual antagonism between two TFs is apparent across development in multiple lineages (Graf and Enver, 2009). For example cross antagonism between TFs GATA1 and PU.1 drives specification of erythroid-megakaryocytic versus monocytic identity in hematopoietic differentiation (Iwasaki and Akashi, 2007), the antagonism between PTF1A and NKX6.1 in pancreatic differentiation drives “tip-trunk” segregation (Schaffer et al., 2010), and mutual inhibition between OCT4 and CDX2 is believed to underlie TE versus ICM specification (Dietrich and Hiiragi, 2007; Niwa et al., 2005). It is possible that mechanisms and network motifs similar to the one we describe here are relevant in some of these contexts to safeguard the dynamic nature of early differentiation and possibly explain why some progenitor cells are readily expandable as stem cells *ex vivo*.

The pluripotency network has been one of the most extensively studied models for *ex vivo* expansion. Based on large volumes of ChIP-seq and interaction studies (van den Berg et al., 2010; Chen et al., 2008; Gagliardi et al., 2013; Kim et al., 2008; Marson et al., 2008; Masui et al., 2007; Rodda et al., 2005; Tomioka et al., 2002), the network is thought to act in a concerted feed forward loop to support a highly stable state (Jaenisch and Young, 2008; Silva and Smith, 2008; Young, 2011). As the pluripotent state exists only transiently *in vivo* (Morgani et al., 2017), this begs the question of how such a strong network has emerged with evolution? A highly cooperative state would be expected to respond to changes in TF concentration in a non-linear way, causing rapid commitment to differentiation. However, the network motif that we propose for NANOG and ESRRB, where expression of both factors supports pluripotency, but where ESRRB on its own enables lineage priming, supports the progression of priming and commitment in the dynamic context of *in vivo* development while also providing resistance to fluctuations in *Nanog* gene expression (Chambers et al., 2007; Kalmar et al., 2009; MacArthur et al., 2012; Singh et al., 2007) in the stable context of *in vitro* culture.

### Limitations of study

Because our system (NΔ, NΔ:E) is based on stable ESCs lines, we lack resolution in time and cannot dissect the order of events between ESRRB expression and the upregulation of a PrE-like identity.

In addition, future work *in vivo* investigating whether ESRRB is indeed important for the observed plasticity in cell identity during Epi PrE lineage segregation is needed to gauge the significance of this mechanism. These experiments will require either maternal-zygotic mutants, *in vivo* degradation systems, double mutants for *Esrrb* and *Nr5a2* or good small antagonists of these factors.

## Acknowledgments

We thank N. Festuccia for *Esrrb*Δ ESCs and the Esrrb-tdT targeting construct, all members of the Brickman laboratory for continuous critical discussion, specifically J.A.R. Herrera for bioinformatics advice and pipeline development and M. Rothová for technical expertise and support on MARS-seq. We thank H. Neil, M. Michaut and the reNEW Genomics Platform for exquisite assistance, support and use of instruments, L. Mariani for assistance with mesoderm differentiation, J. Zylicz for TSCs, K. Stewart-Morgan for advice on ATAC protocols and the manuscript, G. dela Cruz and P. van Dieken for technical support and advice on FACS, J. M. Bulkescher for microscopy support, R. Bone and M. Linneberg-Agerholm for in house processing/analysis of published ATAC-seq and post-implantation single cell RNA-seq respectively (Pijuan-Sala et al., 2019; Wu et al., 2016). The work was funded by grants from the Danish National Research Foundation (DNRF116) and the Novo Nordisk Foundation (NNF17CC0027852 and NNF21CC0073729).

## Author contributions

TEK, WBH, AT and JMB conceived the study. TEK, WBH and JMB designed and interpreted experiments. TEK performed all experiments, except the Esrrb truncation/mutant assessment, which was done by ML. TEK carried out the bioinformatics analysis, except processing of single cell sequencing data and in house analysis of Nowotschin et al., 2019, which was performed by MP. TEK, AT and AVN wrote and analyzed the ODE model. TEK, AT and JMB wrote the manuscript with input from all other authors.

## Methods

### ESC culture and differentiation

All ESC lines were maintained in SL (Serum/LIF) medium: GMEM (Sigma) supplemented with 10% fetal bovine serum (FBS) (Gibco), 100 μM 2-mercaptoethanol (Sigma), 1× MEM non-essential amino acids, 2 mM l-glutamine, 1 mM sodium pyruvate (all from Gibco), 1,000 U ml−1 leukemia inhibitory factor (LIF) (made in house) and passaged with Trypsin. For serum free culture, we used 2iL (2 inhibitors + LIF (Ying et al., 2008)): Neurobasal medium and DMEM:F12 (Gibco) supplemented with N2 (made in house), B27 (Gibco), 3 μM Gsk3i (Chir99021: Axon Medchem) and 1μM MEKi (PD0325901: Sigma) and LIF, and passaged with Accutase (Sigma). Selection for transgenes or targeted cell lines was done using puromycin (1ug/mL, Sigma), neomycin/G418 (100ug/mL, Geneticin), or hygromycin (125ug/mL, Roche). Neural monolayer differentiation was performed as outlined in (Ying et al., 2003); primitive endoderm differentiation was performed as outlined in (Anderson et al., 2017), TSCs were cultured as described in (Tanaka et al., 1998), and mesoderm differentiation was performed as described in (Mariani et al., 2021). Cell lines used in this study are listed in Table S15.

### Immunofluorescent staining, imaging, and analysis

Cells were washed and fixed in 4% formaldehyde (FA) at 37°C for 15 min (Fisher Scientific, PI-28906), permeabilized in ice cold methanol at −20°C, 10min and blocked in 5% donkey serum and 0.3% Triton 1hr at room temperature or overnight at 4°C. Primary antibody incubation was done overnight at 4°C in 1% BSA, 0.3% Triton in PBS, subsequently incubated with the appropriate fluorophore-conjugated secondary (AlexaFluor, molecular probes), DAPI stained and visualized on a confocal Leica TCS SP8 microscope. The widefield Zeiss Axio Observer microscope was used to image colonies for the clonal analysis in Figure 3F. See Table S16 for a list of antibodies and concentrations used. Analysis was carried out using open-source software FIJI (ImageJ) and Python3, scimage (see github repository).

### Flow cytometry analysis and cell sorting

Cells were dissociated with Accutase and incubated with the appropriate conjugated antibodies in 10% FBS-PBS for 20 min, washed extensively and analyzed on an LSRFortessa (BD Biosciences) or analyzed and sorted on the Aria III (BD) or SH800 (Sony) on highest purity for the respective machines. Dead cells were excluded based on DAPI inclusion. A gating strategy example can be found in Figure S1H. If needed, compensation was carried out using single color controls. Antibodies used is listed in Table S16.

### Quantitative PCR with reverse transcription

Total RNA was collected using the RNeasy Mini Kit (Qiagen). Genomic DNA was eliminated by DNase treatment (Qiagen) and 1μg of total RNA was used for first strand synthesis using SuperScript III reverse transcriptase according to the manufacturer’s instructions. cDNA corresponding to 10 ng total RNA was used for RT–qPCR analysis using the Roche LC480 and target amplification was detected with the Universal Probe Library system. Relative concentrations were calculated, and all concentrations were normalized to the geometric mean of 3 housekeeping genes (GAPDH, TBP and PBGD). See Table S17 for a list of primers and probes used.

### Cell line generation

The *pPyCAG-IP* vector (Chambers et al., 2003) was targeted with cDNA for *Esrrb* (short or long isoform) or *Nanog* for generation of the stable lines analyzed in Figures 2B, C. The titration cell line NΔ:cE:iN, was generated by cloning the *Esrrb* and *Nanog* constructs schematized in Figure 3A into the *pPyCAG* and *tetON* (Hamilton et al., 2019) vectors respectively. To generate stable lines constructs were linearized and introduced into *NanogΔ* cells by electroporation followed drug selection and clonal expansion. Chosen clones were carefully selected based on physiological levels of expression as measured by western blot against control WT cells in SL and 2iL. The cell line thus could support expression of either only ESRRB or both ESRRB and NANOG, in the same clonal background and at physiological levels comparable to WT mESCs. Double protein fusion reporter cell line for ESRRB and NANOG was generated in an E14JU WT background, starting at passage 9, and the methodology was inspired by (Festuccia et al., 2018a; Sokolik et al., 2015). First *Nanog* was targeted using CRISPR-Cas9, and the WT allele was sequenced to ensure the PAM site was intact. Second, we targeted *Esrrb* using homologous recombination. Prior to clonal expansion, targeted cells were screened by flow cytometry, no additional selection was used. Cell lines used and generated in this study are listed in Table S15.

### Southern blot analysis

*Esrrb* and *Nanog* targeting was validated according to the strategies presented in (Festuccia et al., 2018a; Sokolik et al., 2015). Southern blot analysis was performed as described in (Southern, 2006) on 5-10ug genomic DNA. [a-32P] dCTP was used to label specific probes, sequences listed in Table S18), and membranes were developed on high resolution Typhoon FLA 7000 biomolecular imager.

### Karyotype characterization

Karyotyping was performed in house on expanding cells using 0.1ug/mL demecolcine/ KaryoMAX (Gibco). After incubation for ∼1hr at 37°C, dividing cells were harvested, washed, and incubated in 75mM KCl for 6min at room temperature before fixation and permeabilization in 75% MeOH, 25% acetic acid over night at 4°C. Cell preparations were splashed onto SuperFrost adhesion slides (Thermofisher), stained in 10% Giemsa, mounted, and imaged using a confocal microscope, Zeiss 780.

### Half-life measurements

Half-lives were assessed by treating relevant cell lines with the translational elongation inhibitor cycloheximide (Sigma-Aldrich 01810-1G) at 20ug/mL for a time course of 3hrs. Cell lysates were processed for western blot analysis as described. Normalization was done to total protein as measured by Coomassie staining. Data was fitting with an exponential regression model (y=ba^x^), y specified to 0.5 and T_½_ calculated as log(0.5/b)/log(a), using the coefficient from the fit.

### Western blot analysis and protein quantification

Protein extracts were prepared by sonication of cell lysates in 4% SDS, 20% glycerol and 120mM Tris-HCl pH 7.4 and quantified on a NanoDrop2000 spectrometer based on 280nm absorption. 20-40ug protein per sample was denatured by heat and DTT (100mM), resolved by electrophoresis in 4-12% Bis-Tris gradient SDS-polyacrylamide gels (NuPAGE, Thermofisher) and transferred to nitrocellulose membranes (GE Healthcare) in 25mM Tris, 192mM glycine and 20% MeOH. Membranes were blocked in 150mM NaCl, 10mM Tris, 0.1% Tween (TBST) with 10% non-fat dried milk and probed with respective primary antibodies in TBST with 5% BSA (Sigma Aldrich) over night at 4°C. Primary antibodies were detected using the appropriate fluorescently conjugated secondary antibodies (Alexa Flour, Molecular Probes), visualized with Chemidoc MP (BioRad) and quantified using FIJI (ImageJ). Total protein staining of the gels by Coomassie and membrane staining of H3 was used as loading control and for normalization. Antibodies used are listed in Table S16.

### Bulk RNA-sequencing and raw data processing

The experiment was performed on the flow cytometry sorted PECAM+PDGFRA-(ESC/Epi-like) fraction of NanogΔ and NanogΔ:cEsrrb:iNanog, with either induced or not induced NANOG expression, from stable cultures in SL. Total RNA was purified by standard methods and rRNA depleted using the NEBNext rRNA depletion kit (NEB, as per the manufacturer’s instructions). RNA-seq libraries were prepared on-bead using the NEBNext Ultra kit as per manufacturer’s instructions and subsequently sequenced using a Next-Seq 500 Sequencer (Illumina). Raw reads were processed with bcl2fastq (v 2.19.1) and STAR (v 2.5.3a) was used to map sequencing reads and to generate the count table (Dobin et al., 2013). Data processing was performed using Computerome, the National Life Science Supercomputer at DTU (www.computerome.dk). Principal component analysis and differential expression analysis were performed using the DESeq2 (Love et al., 2014) and factoextra (Kassambara and Mundt, 2020) packages in R, significance was defined as abs(log2FC) > 1 and adjusted p < 0.01 (Wald’s test p-values were corrected using the Benjamini-Hochberg procedure). Gene overlap analysis was performed in R, using the GeneOverlap package (Shen, 2022), that calculates overlap between all pairs from two gene lists and determines p-value and odds ratio in comparison to a genomic background by a one tailed (greater than) Fisher’s exact test.

### ATAC-sequencing and analysis

The experiment was performed on the exact same samples (divided into two) as the RNA-seq (see above). Nuclei preparation and transposase treatment was carried out as described in (Stewart-Morgan et al., 2019), following this libraries were prepared on-bead using the NEBNext Ultra kit as per manufacturer’s instructions and subsequently sequenced using a Next-Seq 500 Sequencer (Illumina) and paired end 150bp sequencing. Raw reads were processed with bcl2fastq (v 2.19.1), adapters were trimmed with cutadapt (v 2.2.0), reads were mapped to mm10 using bowtie2 (v 2.2.5) (-X 1500 –no-mixed –no-discordant) (Langmead and Salzberg, 2012), chrM and duplicate reads were removed, quality filtering was performed with samtools (v 1.4.1) (-b -f3 -F4 -F8 -q7) and reads in blacklisted regions from (Buenrostro et al., 2013; The ENCODE Project Consortium, 2012) were removed. RPKM normalized bigwigs were generated from filtered bam files for visualization using deepTools bamCoverage (--binSize 5 --normalizeUsingRPKM) (Ramírez et al., 2016). Peaks were called using macs2 (v 2.2.0) (-f BAMPE, -q 0.001), consensus peaks per condition were defined as peaks present in all 3 biological replicates, and these were combined to generate a count table for downstream analysis (reads within defined peaks were counted using bedtools, v 2.26.0, coverage). Principle component analysis and differential enrichment analysis were performed using the DESeq2 (Love et al., 2014) and factoextra (Kassambara and Mundt, 2020) packages in R, significance was defined as abs(log2FC) > 1 and adjusted p < 0.01 (Wald’s test p-values corrected by using the Benjamini-Hochberg procedure). Condition specific enriched peaks were used for 1) motif analysis with HOMER (findMotifsGenome.pl, - size given, background: all consensus peaks) (Heinz et al., 2010) and 2) annotation with HOMER (annotatePeaks.pl, default parameters) for GeneOverlap analysis as described for bulk RNA-seq.

### ChIP-sequencing and analysis

Chromatin Immunoprecipitation and data processing was performed for ESRRB as described in (Hamilton et al., 2019), using double fixation, mapped to mm10 with bowtie2 (Langmead and Salzberg, 2012) and quality filtered with samtools (v 1.4.1) flags (-b -F4 -q13). Normalized bigwigs for visualization were generated with bamCoverage (v 4.4.0) using binsize 5 and RPKM. Peaks were called using macs2 (v 2.2.0) against condition and replicate specific input samples specifying *q* ≤ 0.01, consensus peaks per condition were defined as peaks present in all 3 biological replicates, and these were combined to generate a count table for downstream analysis (reads within defined peaks were counted using bedtools, v 2.26.0, coverage). Differential enrichment analysis was performed using the DESeq2 (Love et al., 2014) package in R, and significance was defined as abs(log2FC) > 1 and adjusted p < 0.05 (Wald’s test p-values corrected by using the Benjamini-Hochberg procedure). Condition specific enriched peaks were used for 1) motif analysis with HOMER (findMotifsGenome.pl, -size 7-10) (Heinz et al., 2010), 2) annotation with HOMER (annotatePeaks.pl, default parameters) for GeneOverlap analysis as described for bulk RNA-seq and 3) binding of GATA6 and NANOG were assessed with deepTools (Ramírez et al., 2016) at these specific regions upon processing of previously published ChIP-seq for these factors (Marson et al., 2008; Wamaitha et al., 2015), using the in house pipeline described here.

### Single cell MARS-seq2

MARS-seq2 was performed as described in (Rothová et al., 2022), with modifications according to (Keren-Shaul et al., 2019), on three biological replicates of the *NanogΔ* cell line and three biological replicates of the NΔ:cE:iN cell line, with constitutive ESRRB expression but without NANOG induction. Even proportions of the chosen populations were sorted onto plates before library preparation. Raw sequencing reads were converted to paired-end fastq format files using bcl2fastq (v 2.19.1). The pre-processing was done using the nf-core/marsseq pipeline (Proks *in prep* 2022, github: https://github.com/nf-core/marsseq) with the following command: nextflow run . -c dangrpufl01.conFigure –genomes_base ./references –input ./design.csv. Raw counts were converted to anndata using scanpy (v 1.8.2). Cells were filtered out based on the following thresholds (800 < #UMIs < 30,000 and 400 < # genes < 5,000). Additionally, we removed empty cells (“Zero” in metadata) and discarded ERCC (External RNA controls consortium) spike-in “genes” and ribosomal genes resulting in 264 cells and 53,546 genes. The counts were CPM normalized and log transformed. The PCA was computed on top 2,000 variable genes followed by UMAP dimension reduction with default settings. Clustering was performed using Louvain clustering with 0.3 resolution identifying 3 cell populations and marker genes for each cluster were extracted with thresholds log2FC > 0.5 and adj. p-val < 0.05. GeneOverlap analysis was done as described for bulk RNA-seq, and CAT analysis was performed as described in (Rothová et al., 2022) with default settings. We used normalized and log transformed counts as input for both datasets (see https://github.com/brickmanlab/CAT for the code).

### In house processing and velocity estimation for external single cell sequencing data

To estimate velocity on pre-implantation blastocyst single cell sequencing (Nowotschin et al., 2019), we first downloaded fastq files from GSE123046 using ffq (v 0.0.4) (Gálvez-Merchán et al., 2022) with the following command: ffq -t GSE -o kat.json GSE123046, and only E3.5 and E4.5 data was processed further. Next using STAR (v2.7.3a) we generated genome references to mm10 (Ensembl release 98) and we used STARSolo to generate raw, spliced and unspliced counts (Dobin et al., 2013). We used 737K-august-2016 10X whitelist to identify correct cell barcodes (https://github.com/10XGenomics/cellranger/blob/master/lib/python/cellranger/barcodes/737K-august-2016.txt). We concatenated spliced and unspliced count matrices with the original count matrix provided in the publication (partitioned on E3.5 and E4.5 stages). Downstream analysis was performed using scvelo (v0.2.4) with default settings using dynamical modeling (Bergen et al., 2020). Using Mann Whitney U Test (Wilcoxon rank sum test) we identified differentially expressed genes (markers) with the following thresholds: Log2FC > 1 and adj. p-val < 0.05. For the post-implantation cell type marker lists, the published count matrix was downloaded (Pijuan-Sala et al., 2019) and used to extract marker gene lists with Seurat, R (FindAllMarkers, default settings, filtering on adjusted p-value < 0.05) for use in GeneOverlap analysis.

### ODE modeling and stochastic simulations

To allow comparison between a simple GRN consisting of GATA6, NANOG and FGF to the GRN with ESRRB, we first wrote a 2-variable differential equation system for the simple GRN using multiplicative logic and cooperative interactions represented by Hill coefficients (for a description of all parameters used see Table S19):

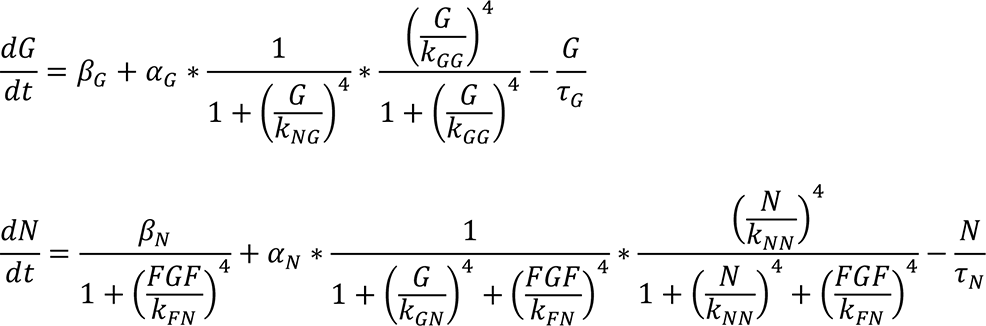

All programming was done using Python3. We tuned the parameters to support a model that replicates the core principles of FGF-mediated Epi-PrE segregation, namely that FGF inhibition gives all Epi identity, overstimulation gives all PrE identity and endogenous FGF allows the two populations to specify side by side (Ambrosetti et al., 1997; Chazaud et al., 2006; Yamanaka et al., 2010). We then extended the model to include ESRRB, assuming that NANOG, when bound to ESRRB, inactivates ESRRB’s ability to activate GATA6:

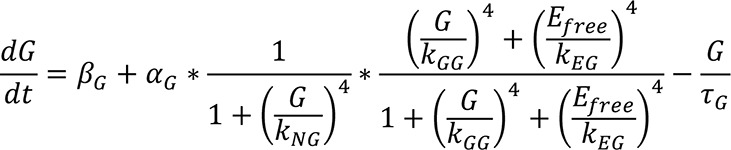

Where 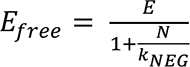 is calculated from mass conservation

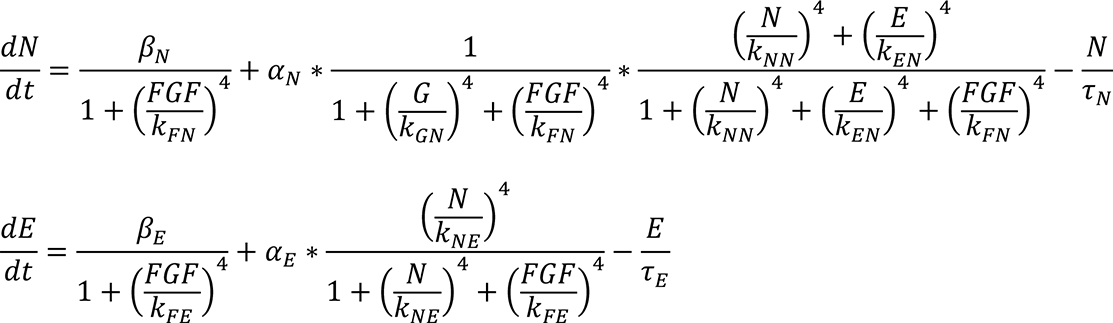

We accessed bifurcation dynamics by numerical integration of trajectories from 2000 randomly chosen initial conditions (ICs) in the range [0.01,17] for G, N and E setting t_max_=200 using scipy.integrate solve_ivp. Phase space diagrams were made using matplotlip streamplot. Time difference between ICs reaching steady state for GNF and GNFE was calculated for 2197 specific ICs in the GATA6^low^-NANOG^low^ region. These were specified by drawing from a log2-uniform distribution between 0 and 5.6, as ICs for both NANOG and GATA6, however restricted to ICs that fulfil IC_GATA6_ + IC_NANOG_ < 5. The ESRRB ICs were calculated from these, by setting ESRRB to steady state (IC values for NANOG and GATA6 are depicted in Figures S7B and C, left panel). Time to reach steady state was then quantified by identifying the time at which the slope of the concentration changes of GATA6 or NANOG along time stagnates and is close to 0.

To simulate changes in the environment for individual cells of the embryo, we reformulated the deterministic ODE model as an event-based stochastic model and implemented it using Gillespie algorithm (Gillespie, 1976). Briefly, each of the terms in the ODEs are interpreted as reaction rates for the reactions that result in either increase or decrease number of molecules by 1. The algorithm consists of two major steps. First, given the numbers of molecules, we calculate all possible reaction rates *rate_i_*, for each reaction *i* we find the times until next event: 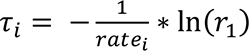. Second, we perform the event corresponding to the reaction with the smallest *r*_i_, and advanced the simulation time by *r*_i_. The two steps are repeated until simulation time reaches a predetermined limit, typically selected long enough for system to reach steady state. We used the specified 2197 GATA6^low^-NANOG^low^ ICs defined above and for all ICs compared the endpoint from the stochastic systems, with that of the deterministic systems and calculated the probability of shifting endpoint when exposed to noise for both GNF and GNFE (Figures 6H and S7D, t=0). To simulate environment changes across lineage specification, we used the deterministic system to propagate IC positioning across time (Figures S7B and C) and then started the stochastic simulations from successively later timepoints (Figures 6H and S7D, t=1 and t=2). See Github repository for code used.

### Data availability

GEO reviewer access token: odclwumahrgrzyf

GEO link: https://www.ncbi.nlm.nih.gov/geo/query/acc.cgi?acc=GSE207565

Main github repository: https://github.com/brickmanlab/knudsen_et_al_2022

## Supplementary information

### Supplementary Figures and Legends S1-7

**Supplementary Figure 1, related to Figure 1.**
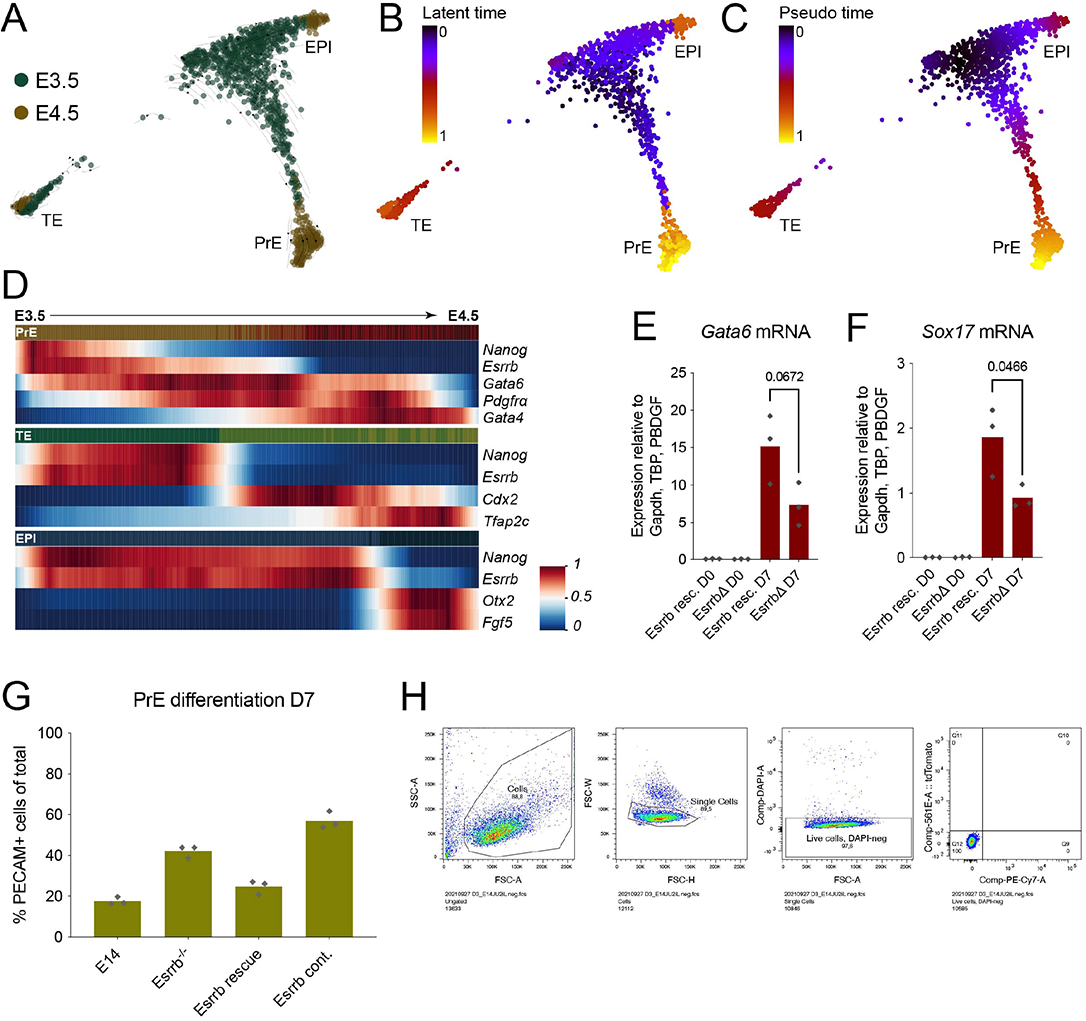
A role for Esrrb in PrE differentiation. **(A)** tSNE embedding of mouse blastocyst single cell sequencing at E3.5 and E4.5, timepoint of origin is labelled (Nowotschin et al. 2019). Projected RNA-velocity was generated using scVelo (Bergen et al. 2020). **(B)** scVelo estimated latent time projected on tSNE embedding as in A. **(C)** Seurat estimated pseudotime projected on tSNE embedding as in A. **(D)** Heatmaps of scaled expression of indicated genes along pseudotime (colored bar at the top of each subplot, colors as in Figure 1A) for PrE (top), TE (middle) and Epi (bottom). **(E-F)** RT-qPCR for the PrE markers, *Gata6* and *Sox17*, expression at D7 of *in vitro* PrE differentiation for the indicated genotypes/ conditions as explained in Figure 1E (unpaired t-test was used for statistical analysis, n = 3). **(G)** Flow cytometry analysis of the ESC marker, PECAM, expression at D7 of *in vitro* PrE differentiation for the indicated genotypes/ conditions as explained in Figure 1E (n = 3, biological clones). **(H)** Flow Cytometry gating strategy applied throughout this study.

**Supplementary Figure 2, related to Figure 1.**
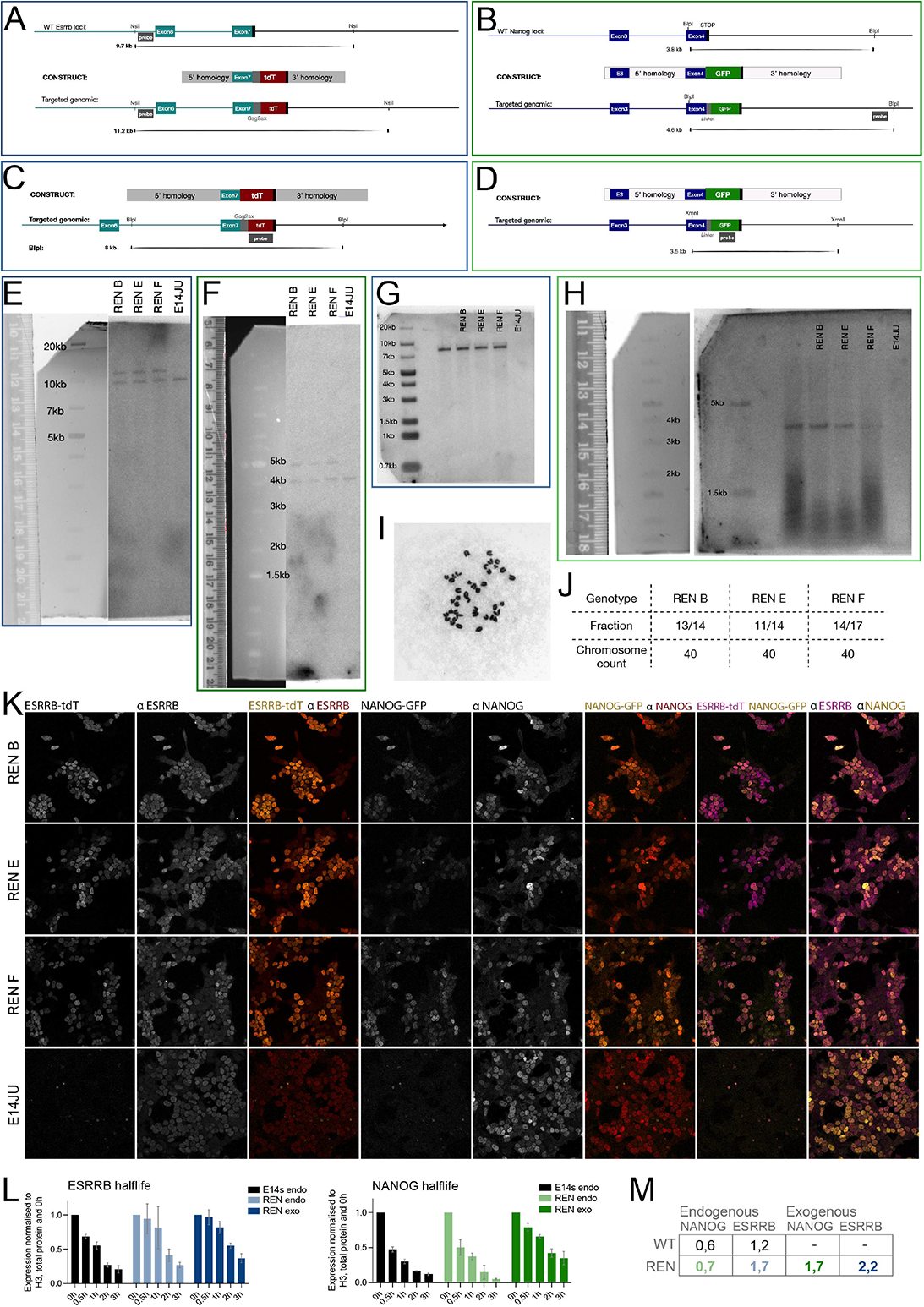
Generation of a double reporter for ESRRB and NANOG (REN) **(A-B)** Southern strategy for validation of targeting tdT to the C-terminal of *Esrrb* (A) and eGFP to the C-terminal of *Nanog* (B), restriction enzyme digests cut outside of the construct homology arms and probes were designed with homology to an endogenous genomic region either 5’ (*Esrrb*) or 3’ (*Nanog*) of the targeting construct. Predicted sizes: WT *Esrrb* locus ∼9.7kB, targeted *Esrrb* locus ∼11.2kB, WT *Nanog* locus ∼3.8kB, targeted *Nanog* locus ∼4.6kB. **(C-D)** Southern strategy for assessing multiple integration sites of tdT (C) or eGFP (D), restriction enzyme digests cut inside of the construct homology arms and probes were designed with homology to either tdT or GFP. The predicted size for targeted integration of tdT and eGFP was 8kB and 3.5kB respectively. **(E-F)** Southern blots corresponding to A and B respectively, validating heterozygote targeting of both the *Nanog* and *Esrrb* loci of REN clones compared to the parent, WT E14JU cell line. **(G-H)** Southern blots corresponding to C and D respectively, validating that tdT and eGFP was only integrated in the targeted loci of REN clones compared, with no integration in parent, WT E14JU cell line. **(I-J)** Karyotype characterisation of REN clones showing a minimum of 78% natural karyotype/clone. **(K)** Immunofluorescent imaging of REN clones and parent, WT E14JU, comparing ESRRB and NANOG expression by staining with the respective antibodies to the tdT/ GFP signal produced by the targeted fusion proteins. **(L)** Western blot quantification of half-life assay on REN clones and WT, E14JU cells, comparing the half-life of the endogenous and fusion proteins (data are mean ±SD, n = 3). Significance was tested using an unpaired two-tailed t-test and no comparisons were found significantly different. **(M)** Table of calculated half-lives from fitting the data in L with an exponential regression model (y=ba^x^) and calculated T_½_=log(0.5/b)/log(a).

**Supplementary Figure 3, related to Figure 1.**
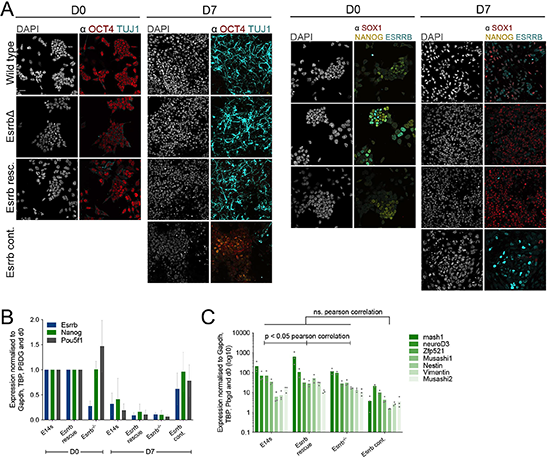
Esrrb is dispensable for neural differentiation. **(A)** Immunofluorescent staining and imaging of indicated markers at D0 and D7 of neural differentiation, across conditions as schematized in Figure 1E (n=3). Scalebar = 30μm, representative for all panels. **(B-C)** RT-qPCR for indicated pluripotency genes (B) and neuronal marker genes (C) at D0 and D7 of neural differentiation, across conditions as in A and schematized in Figure 1E (n=3). Statistical significance was calculated using Pearson correlation.

**Supplementary Figure 4, related to Figure 2.**
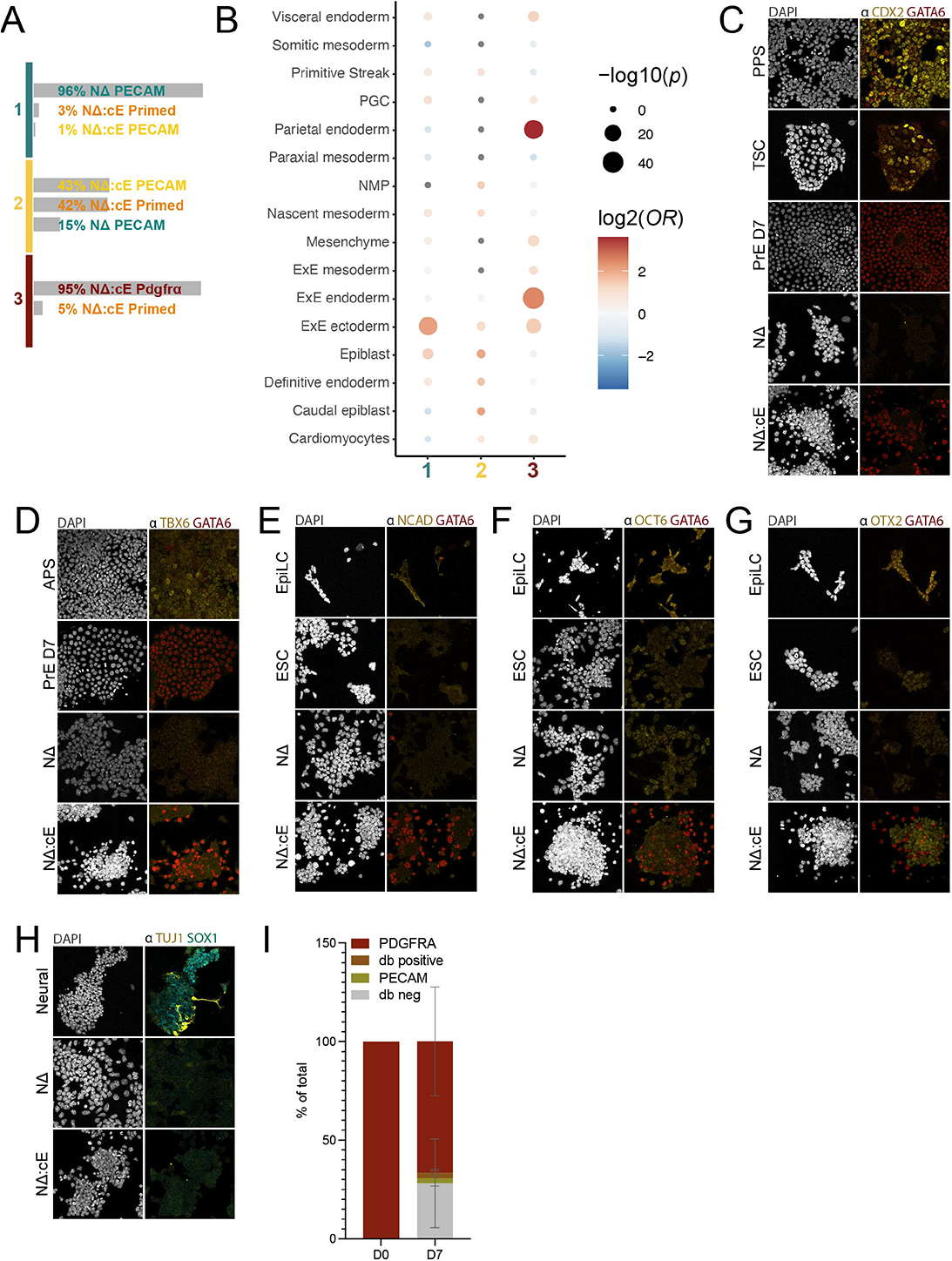
In the absence of Nanog Esrrb support a stable subpopulation of PrE-like cells. **(A)** Overview of cluster composition in the sc-seq data segregated by the sorted populations (see Figures 2D-F). **(B)** GeneOverlap analysis of clusters in Figure 2F against marker lists from *in vivo* single cell RNA seq of post-implantation embryos (Pijuan-Sala et al. 2019). Grey dots indicate Odds Ratio (OR). **(C-H)** Immunofluorescent staining and imaging of indicated cell types and markers. ESC = wild type ESCs in SL conditions, PrE D7 = end point (day 7) of PrE differentiation, TSC = Trophoblast Stem Cells, APS/PPS = 48hr timepoint of anterior/posterior primitive streak differentiation, EpiLC = 48hr timepoint of Epi-Like Cell differentiation, Neural = Day 7 of neural differentiation. **(I)** Wild type PDGFRA+ cells from the end of PrE differentiation (D0) were placed back into ESC (SL) conditions and analyzed by flow cytometry after 7 days in culture (D7). Cells were stained for the PrE marker, PDGFRα, and the ESC marker, PECAM.

**Supplementary Figure 5, related to Figure 4.**
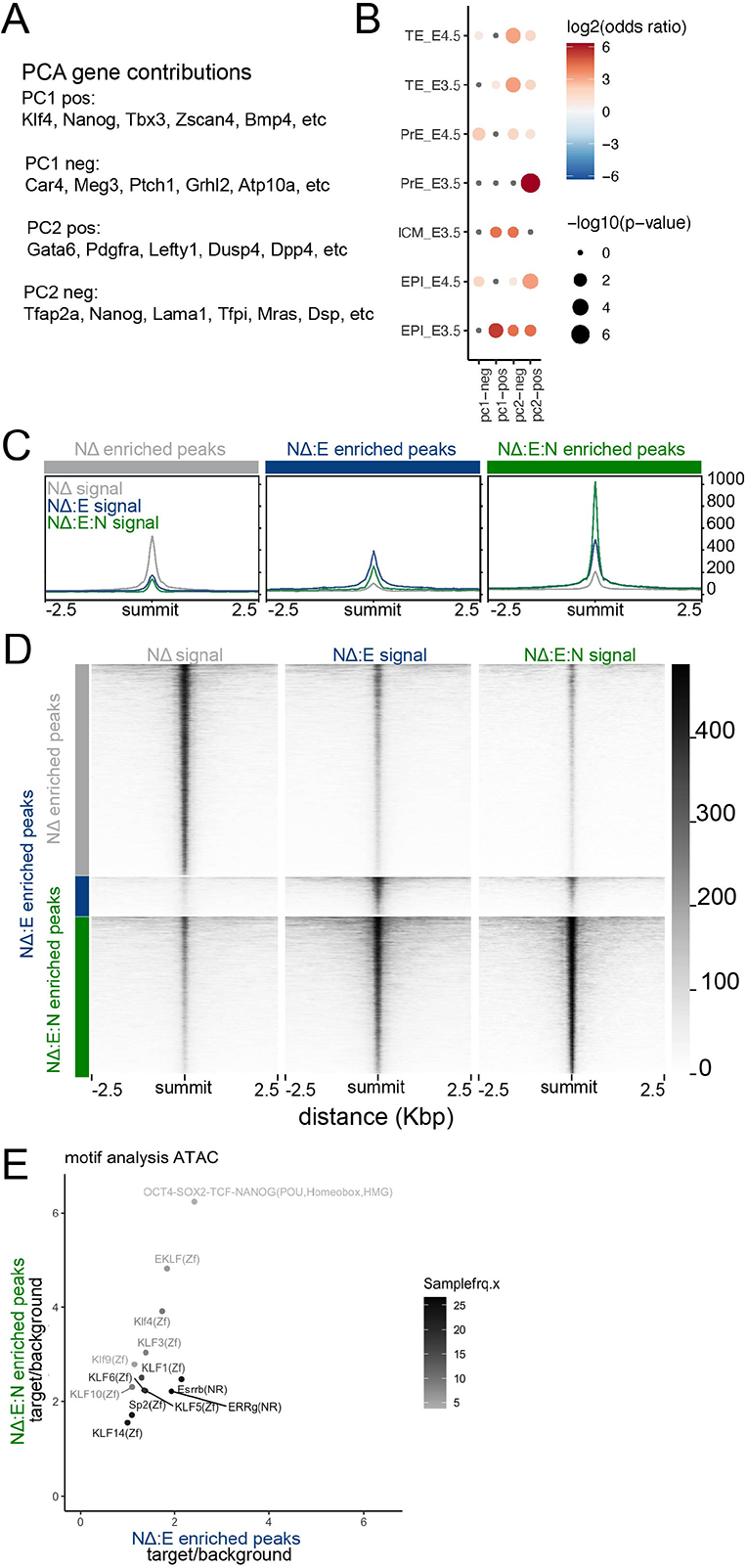
RNA- and ATAC-seq show an ESRRB and NANOG specific responses. **(A)** Genes contributing to the variance represented in Figure 4B. PCs are split into variance that contributes positively (pos) and negatively (neg). **(B)** GeneOverlap analysis of top 50 variance contributing genes for PC1 and 2, positive and negative directions, against marker lists from sc-seq of the pre- implantation blastocyst (Nowotschin et al. 2019). **(C-D)** Signal from ATAC-seq of NanogΔ (NΔ), NanogΔ:Esrrb (NΔ:E) and NanogΔ:Esrrb:Nanog (NΔ:E:N) on the defined specifically enriched peaks for each condition. **(E)** Motif analysis of ATAC peaks specifically enriched when ESRRB was expressed in the absence of NANOG (NΔ:E) vs in the presence of NANOG (NΔ:E:N). Values plotted are motif prevalence in target peaks normalized to the motif prevalence in the genome in general. Gray scale coloring reflects the frequency of the motif in the NΔ:E peaks.

**Supplementary Figure 6, related to Figure 5.**
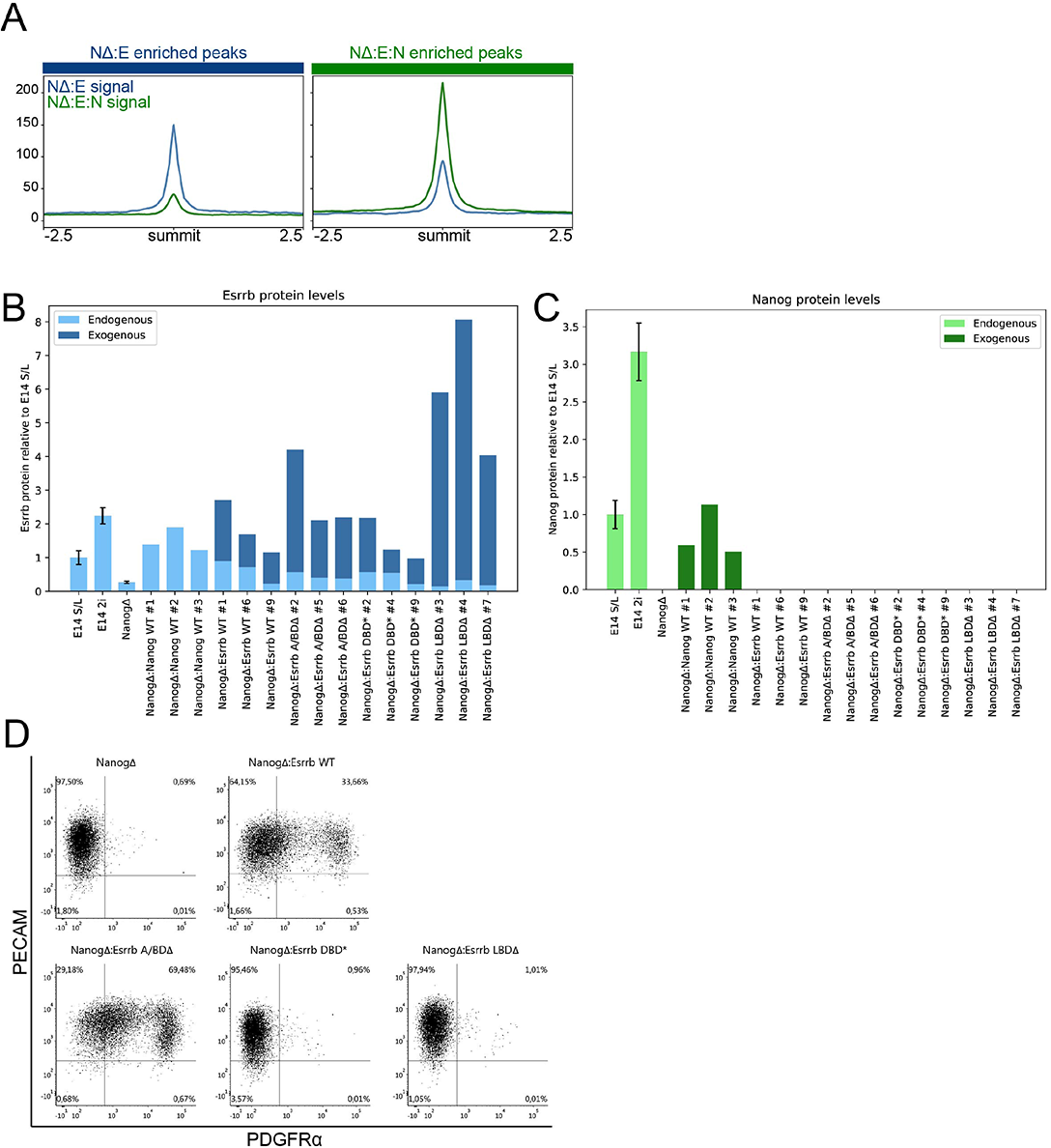
DNA binding by ESRRB is necessary for its PrE affiliated activity. **(A)** Signal from ChIP-seq of ESRRB expressed in the absence (NΔ:E), or presence of NANOG (NΔ:E:N) on the defined specifically enriched peaks for each condition. **(B-C)** Quantification of ESRRB (B) and NANOG (C) expression from western blot analysis after transfection of *NanogΔ* with transgenes encoding for *Nanog* and *Esrrb* as controls, as well as truncations/pointmutations of *Esrrb*: A/BDΔ = N terminal truncation of the transactivating domain, LBDΔ = C-terminal truncation of the ligand binding domain and DBD* = 3 point mutations in the DNA binding domain (Glu142, Lys145 and Lys149 to Gly). WT E14JU cells in SL and 2iL are included as a reference. **(D)** Representative flow cytometry plots from analysis of the PrE marker PDFFRA and the ESC marker PECAM in stable NΔ cell lines expressing the indicated transgenes (n=3, biological clones).

**Supplementary Figure 7, related to Figure 6.**
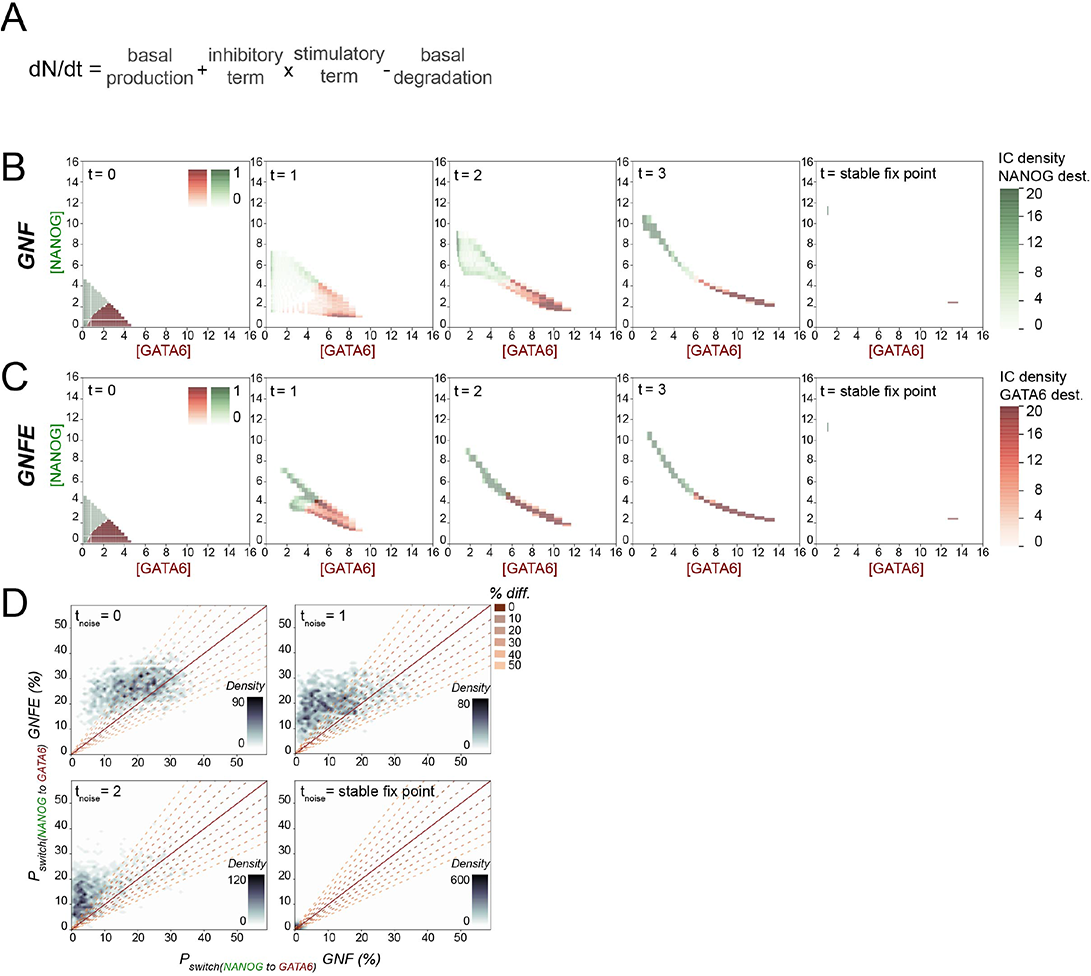
ODE models for PrE-Epi segregation with and without bipartite ESRRB activity. **(A)** Generalised ODE for showing that the change over time in a member for the network, eg Nanog, (dN/dt) is calculated by combining basal production and degration with the stimulatory and inhibitory edges represented in the networks in Figures 6 A, B. **(B-C)** 2d histograms of initial conditions (t=0) and their trajectory through phase space based on the deterministic model (without noise) in the network without (GNF) and with ESRRB (GNFE) at the indicated timepoints. Initial conditions are colored according to their end destination, red represent conditions ending in the GATA6^high^ stable fix point, green represents those ending in the NANOG^high^ fix point. **(D)** Scatter plot of probability of switching fix point destination from NANOG^high^ to GATA6^high^ without (GNF) versus with ESRRB (GNFE). End destination for each of the 2197 initial conditions was defined using the deterministic system (Figures S7B, C). Noise was introduced at 4 distinct timepoints (indicated by t_noise_) during segregation.

### Supplementary Tables 15-19

**Table S15:**
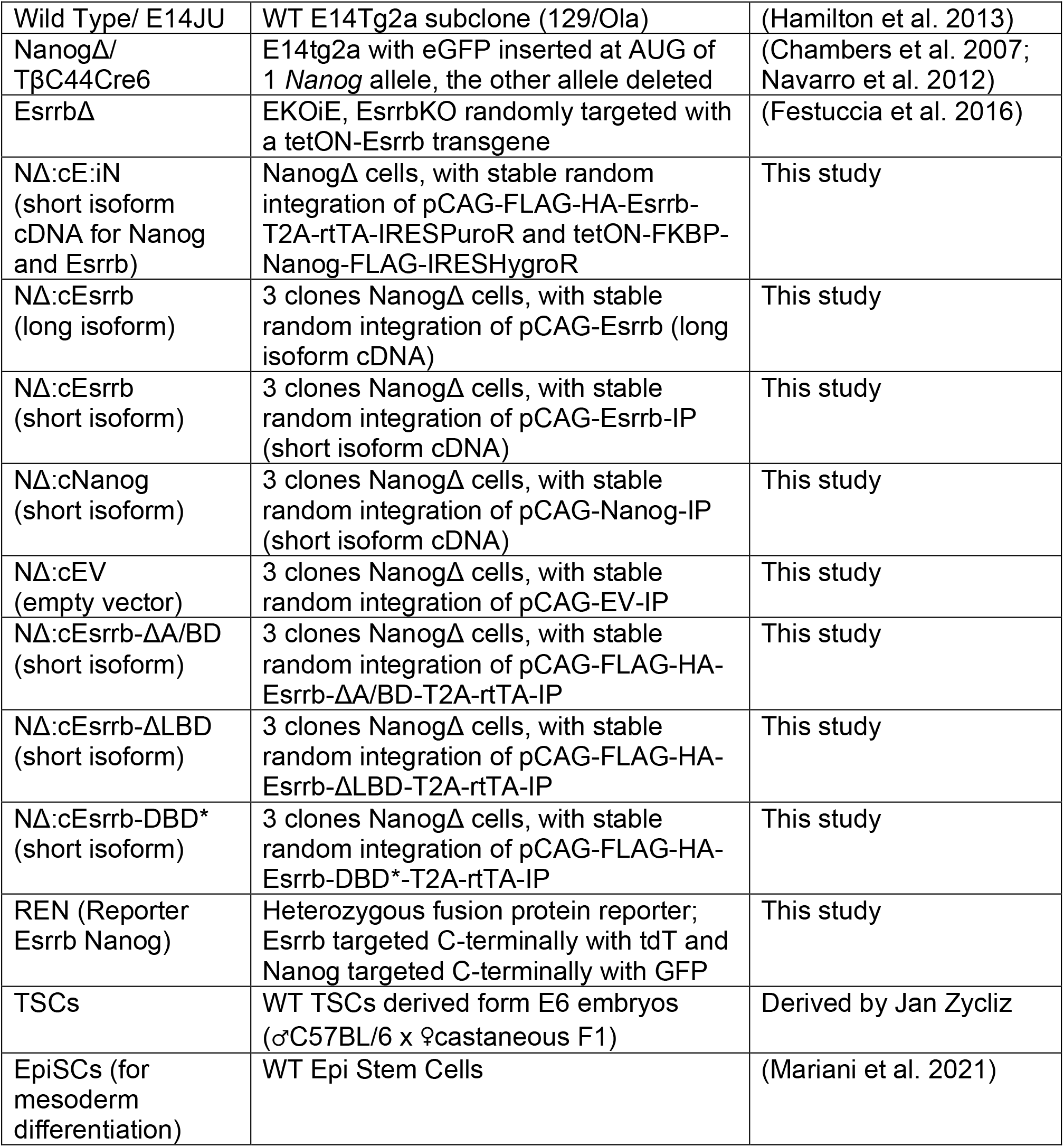
Cell lines.

**Table S16:** Antibodies.

| **Antibody** | **concentration** | **use** | **Supplier and cat. no** |
| --- | --- | --- | --- |
| PDGFRA-PECy7 | 1:200 | FACS | eBioscience, #25-1401-80 |
| PDGFRA-PE | 1:200 | FACS | eBioscience, #12-1401 |
| PECAM-APC | 1:200 | FACS | BD Pharmingen, #551262 |
| GATA6-XP rabbit | 1:1000 | IF | Cell signaling, #5851 |
| NANOG rat | 1:200 | IF | eBioscience, #14-5761 |
| SOX2 goat | 1:100 | IF | R&D, #AF2018 |
| PDGFRA rat | 1:200 | IF | eBioscience, #14-1401 |
| TFAP2C mouse | 1:200 | IF | Santa Cruz, #12762 |
| T-BRA goat | 1:500 | IF | R&D, #AF2085 |
| OCT4 rabbit | 1:200 | IF | Abcam, #ab19857 |
| TUJ1 mouse | 1:1000 | IF | Covance, #mms-435p |
| SOX1 rabbit | 1:200 | IF | Cell Signalling, #4194 |
| CDX2 mouse | 1:250 | IF | BioGenex, #MU392A-UC |
| TBX6 goat | 1:100 | IF | R&D, #AF4744 |
| NCAD mouse | 1:200 | IF | Abcam, #ab98952 |
| OCT6 goat | 1:200 | IF | Santa Cruz, #sc132396 |
| OTX2 goat | 1:200 | IF | R&D, #AF1979 |
| ESRRB mouse | 1:200/1:1000/<br>1ug/5ugChromatin | IF/ WB/<br>ChIP | R&D, #PP-H6705-00 |
| NANOG XP rabbit | 1:1000 | WB | Cell Signaling, #8822 |
| H3 mouse | 1:10.000 | WB | Abcam, #ab171870 |
| *Alexa Fluor, donkey* |  |  |  |
| $\alpha$ rat 647 | 1:2000 | IF | Abcam, #ab150155 |
| $\alpha$ rabbit 568 | 1:2000 | IF | Thermo Fisher Scientific, #A-10042 |
| $\alpha$ mouse 647 | 1:2000 | IF/WB | Thermo Fisher Scientific, #A-31571 |
| $\alpha$ goat 647 | 1:2000 | IF | Thermo Fisher Scientific, #A-21447 |
| $\alpha$ rabbit 488 | 1:2000 | IF/WB | Thermo Fisher Scientific, #A-21206 |

**Table S17:**
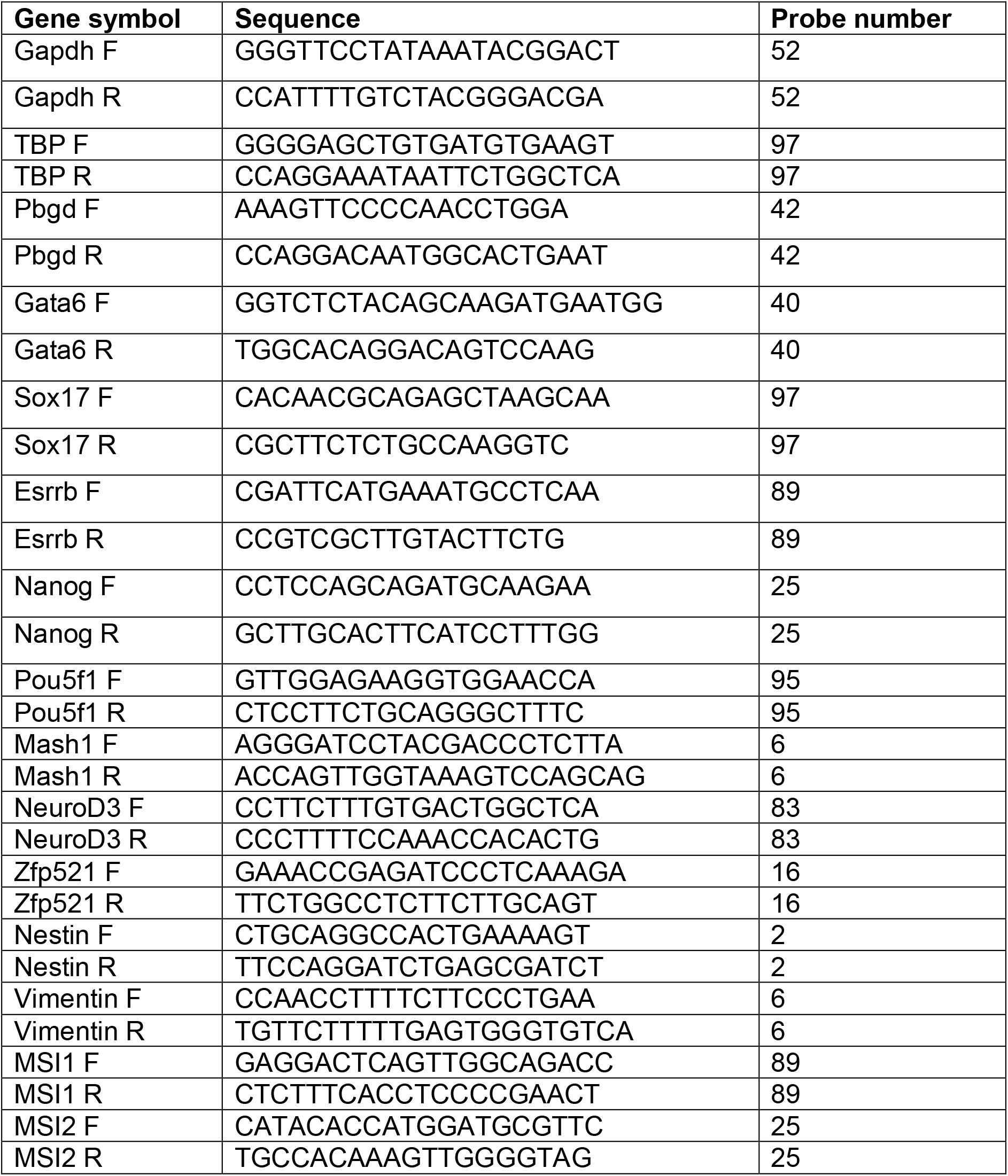
Primers and probes used for RT-qPCR.

**Table S18:**
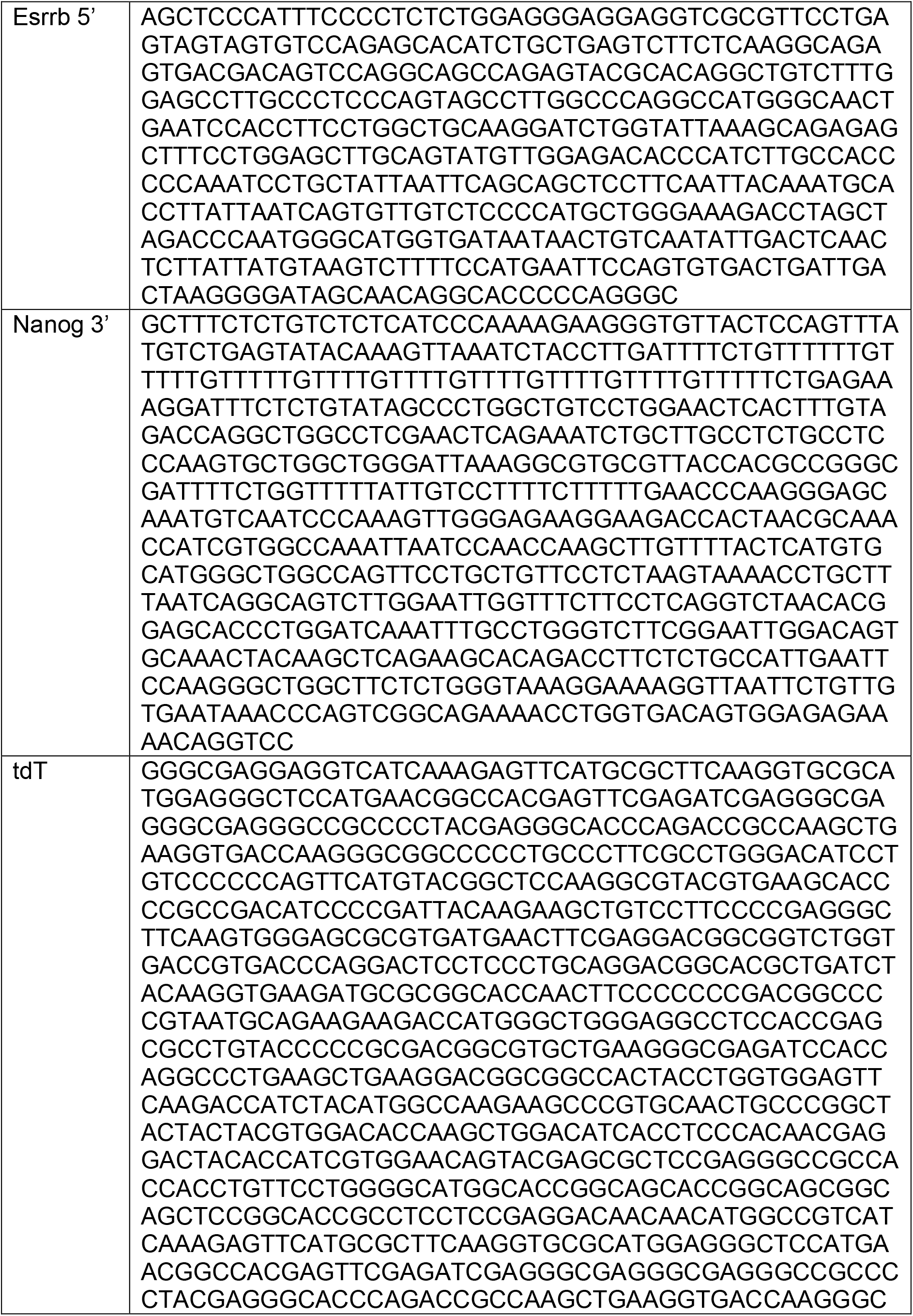
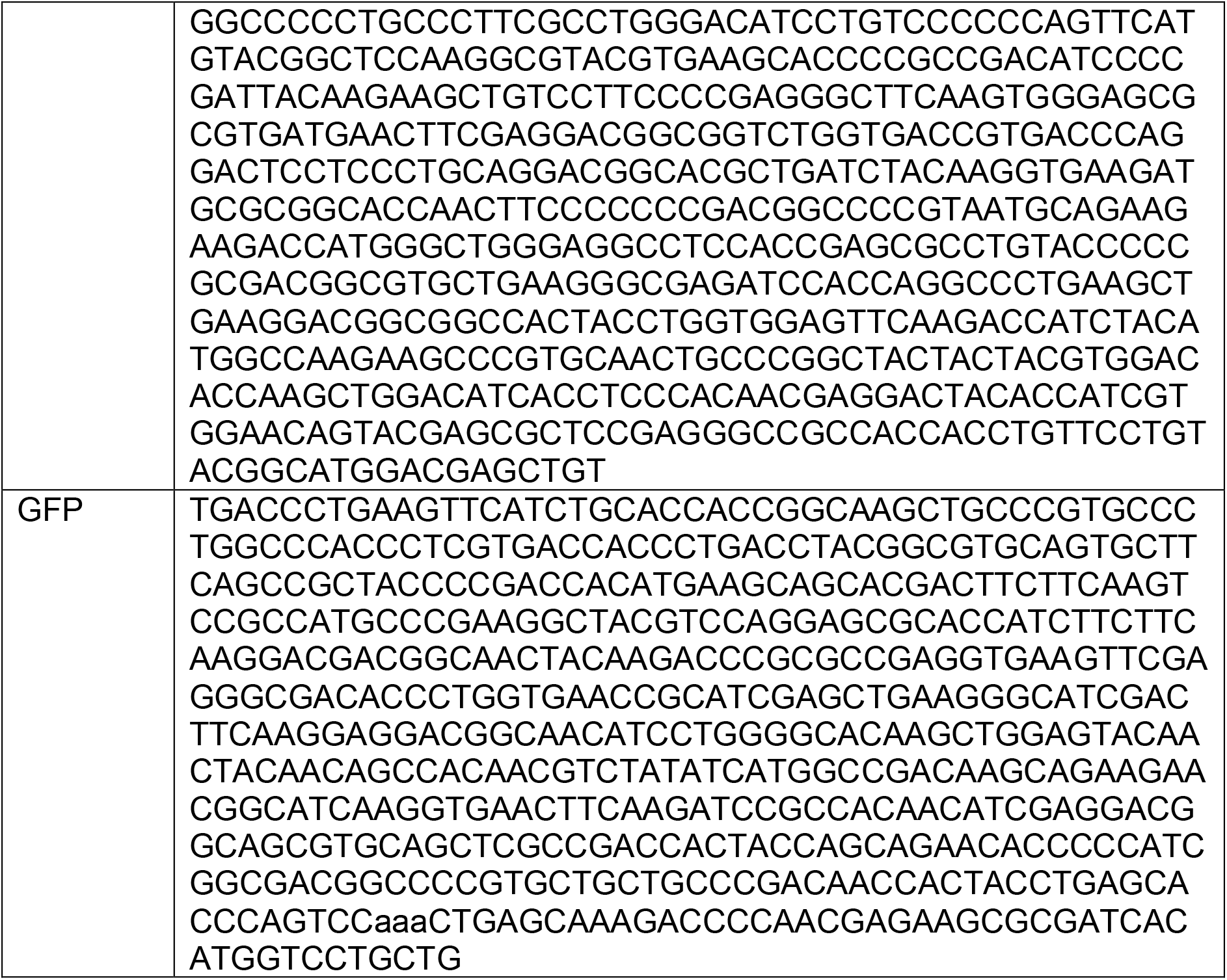
Southern probes.

**Table S19:** Parameters used in ODE models.

| parameter | definition | value | network |
| --- | --- | --- | --- |
| $\beta_N$ | NANOG basal production | 3.5 | GNF/GNFE |
| $\beta_G$ | GATA6 basal production | 1 | GNF/GNFE |
| $\beta_E$ | ESRRB basal production | 1 | GNFE |
| $\alpha_N$ | NANOG weight factor | 14 | GNF/GNFE |
| $\alpha_G$ | GATA6 weight factor | 14 | GNF/GNFE |
| $\alpha_E$ | ESRRB weight factor | 12 | GNFE |
| $\tau_G$ | GATA6 half-life | 1 | GNF/GNFE |
| $\tau_N$ | NANOG half-life | 1 | GNF/GNFE |
| $\tau_E$ | ESRRB half-life | 1 | GNFE |
| $k_{GG}$ | Rate constant for GATA6 autoregulation | 1.2 | GNF/GNFE |
| $k_{NN}$ | Rate constant for NANOG autoregulation | 1.2 | GNF/GNFE |
| $k_{GN}$ | Rate constant for GATA6 inhibiting NANOG | 4.2 | GNF/GNFE |
| $k_{NG}$ | Rate constant for NANOG inhibiting GATA6 | 4.2 | GNF/GNFE |
| $k_{FN}$ | Rate constant for FGF inhibiting NANOG | 1 | GNF/GNFE |
| $k_{FE}$ | Rate constant for FGF inhibiting ESRRB | 3 | GNFE |
| $k_{EN}$ | Rate constant for ESRRB stimulating NANOG | 1.2 | GNFE |
| $k_{NE}$ | Rate constant for NANOG stimulating ESRRB | 1.2 | GNFE |
| $k_{NEG}$ | Rate constant for NANOG inhibiting ESRRB mediated GATA6 stimulation | 1.2 | GNFE |
| $k_{EG}$ | Rate constant for ESRRB stimulating GATA6 | 1.2 | GNFE |
| h | Cooperative interactions, hill coefficient | 4 | GNF/GNFE |
| FGF | FGF concentration | 0.85* | GNF/GNFE |

\*Unless otherwise specified

